# Going green: Recycling transcriptomes to infer evolutionary relationships, gene duplication, gene tree conflict, and patterns of molecular evolution in the Apocynaceae

**DOI:** 10.1101/2025.06.20.660724

**Authors:** Shawn Arreguin, Nathanael Walker-Hale, Madeline R. Casagrande, Savannah Bishop, Eliza Pugacewicz, Mohammed Ramizuddin, Caili Savitzky, Natalia Morales, Arnold Dhan, Beatriz Dybas da Natividade, Valorie Madrid, Cara Simmons, Alexia Diaz, Dena Ahmed, Zoey Ho, Jesenia Nieto Pellecer, Sarahy Diaz, Tesia Mathew, Tony Mirshed, Brett M. M. Simon, Nandrew Thai, Sean Dufault-Hunter, Jarrad Hampton-Marcell, Joseph F. Walker

## Abstract

**Background and Aims:** The flowering plant family Apocynaceae exhibits diverse adaptations with biological and pharmaceutical significance, many of which have been studied with RNA-seq. However, despite the available transcriptomic data, no focused phylotranscriptomic study has been conducted to characterize the patterns of molecular evolution in this group. In this study, we leverage a dataset composed predominately of publicly available transcriptomes to infer relationships within Apocynaceae and explore the molecular processes that have shaped their divergences.

**Methods:** We extracted nuclear, chloroplast, and mitochondrial genes from 47 publicly available and one newly sequenced transcriptome to assemble and infer species relationships across Apocynaceae. Leveraging the gene-rich nuclear phylotranscriptomic data, we inferred molecular dates and investigated the complex history of gene tree conflict, molecular rate shifts, and gene duplications. To investigate the genomic basis of adaptations, we analyzed the inferred 14,838 gene duplications at the base of Apocynaceae for shared functional enrichment of genes related to evolutionary innovations.

**Key Results:** The Apocynaceae topology inferred from our phylotranscriptomic analysis is highly concordant with the current consensus. Notably, a genome-wide acceleration in molecular rate subtends the Ceropegieae tribe. We observed that a decreased time between divergences is associated with a higher rate of gene tree conflict, a pattern especially prevalent across the Apocynaceae’s historically recalcitrant backbone relationships. Furthermore, gene duplications may underlie evolutionary innovations, such as immunity-related gene expansion in the genus *Asclepias* and duplications associated with trichome modifications in the epiphytic *Hoya*.

Finally, we discuss the contentious history of whole-genome duplication (WGD) within the Apocynaceae and emphasize the need for further investigation into the placement of WGDs.

**Conclusions:** Repurposing transcriptomes is a powerful means of accumulating data for novel insights in plant evolution, especially during uncertain funding and cases where budget restrictions exist. Leveraging this gene-dense dataset, we obtained novel insights into the molecular evolution of Apocynaceae and identified areas for future investigations.

## Introduction

The flowering plant family Apocynaceae, commonly referred to as the dogbane family, has a cosmopolitan distribution with species native to all continents except Antarctica (**Bitencourt et al., 2021).** The life history of the group ranges from short-lived herbaceous annuals (e.g., *Conomitra lineari*s) to woody perennials (e.g., *Alstonia scholaris*) (**Endress and Bruyns, 2000**). Several species produce defensive compounds, such as Cardenolides (e.g., *Asclepias syriaca*), latex (e.g., *Apocynum cannabinum*), and Pyrrolizidine Alkaloids (e.g., *Parsonsia alboflavescens*) (**Livshultz et al., 2018a**), which has made the family a source for drug discovery and pharmaceutical research (**Bhadane et al., 2018**). The clade has also been a focus of studies of unique adaptations such as trap flowers (**Heiduk et al., 2010**) and pollen transfer (**Livshultz et al., 2018b**).

Insights into the evolutionary relationships of the Apocynaceae family have come from a single locus (**Sennblad & Bremer, 1996**; **Rapini et al., 2007**), multi-gene (**Bell et al., 2010**), whole plastome (**Fishbein et al., 2018; Wang et al., 2023**) and Angiosperms353 probe datasets (**Antonelli et al., 2021**). These studies have provided a backbone phylogeny for the clade’s five subfamilies. The evolutionary context has helped researchers understand the biogeography and morphological evolution of the groups, approximately 350 genera and 5,000 species (**Fishbein et al., 2018; Bitencourt et al., 2021**). Despite species-rich analyses, large transcriptomic datasets have not been used to interrogate the relationships within the clade.

Phylotranscriptomics, using transcriptomes for phylogenomic analysis, is a cost-effective method approach to plant phylogenetics (**Yang et al., 2017**). Through the thousands of gene trees generated by these datasets, researchers are afforded the rare opportunity to investigate the processes underlying species relationships. Gene and genome duplications (**Smith et al., 2015**; **Yang et al., 2018; Feng et al., 2024**), gene tree conflict (**Maddison, 1997; Rokas et al., 2003**; **Smith et al., 2015**) and signatures of positive selection (**Wang et al., 2019**) are just a few insights provided by phylotranscriptomics. Furthermore, the gene-rich data can be combined with other data types (**Lagou et al., 2024**) to refine our understanding of evolutionary relationships (**Leebens-Mack et al., 2019**).

Phylotranscriptomics relies upon transcriptome data. Therefore, many studies can increase their sampling by repurposing data originally obtained for other transcriptomic studies (**Wong and Peakall, 2022**), such as differential expression (e.g., **Verma et al., 2014**) or genome annotation (e.g., **Park et al., 2014**). Repurposing data provides a means to increase the utility and impact of previous studies. The diverse adaptations found in Apocynaceae have led to a broad range of differential gene expression studies, making the family an ideal candidate for phylotranscriptomics using repurposed data.

This study explores the utility of repurposing differential gene expression data for phylotranscriptomic analysis of Apocynaceae. With this data, we use the most gene-dense analysis to date to investigate the evolution of the dogbane family’s nuclear, chloroplast, and mitochondrial genomes. We explore the group’s propensity for gene/genome duplication, conflict among genes, and inferred shifts in the rate of molecular evolution. Our findings demonstrate that recycling transcriptomic data can yield novel molecular evolutionary insights, demonstrating its value for plant phylotranscriptomics.

## Materials and Methods

### Transcriptome data acquisition and assembly

We extracted and sequenced RNA from silica-dried leaf tissue of *Asclepias viridiflora* left at room temperature for two years, similar to **Ruiz-Vargas et al. (2024)**. The library preparation and sequencing were conducted alongside the samples from **Tyszka et al. (2025)** using the same procedure. To complement this, we downloaded sequencing data for 47 ingroup taxa from the plant family Apocynaceae and eight outgroup taxa from the order Gentianales to broadly cover groups sister to the Apocynaceae (**Supplementary Table 1**).

The raw sequencing data was first processed using the *preprocess* function of the Semblans v1.0.2 package (**Woodcock-Girard et al., 2025**) until the Trimmomatic **(Bolger et al., 2014)** step. In short, Semblans first filtered the reads deemed unfixable based on the Rcorrector v1.0.7 metrics (**Song and Florea, 2015**). The remaining reads were processed using Trimmomatic v0.39 for quality control and adapter removal. The paired Trimmomatic reads were filtered to remove bacterial, protist, fungal, and human contamination using Kraken2 v2.1.2 (**Wood et al., 2019**) with the STD_PF database (https://genome-idx.s3.amazonaws.com/kraken/k2_pluspf_20220607.tar.gz). The chloroplast and mitochondrial reads were extracted from the remaining reads based on the Kakapo v1.0.2 (**Ramanauskas and Igic, 2023**) chloroplast and mitochondrial databases using Kraken2 with a confidence value of “0.1”. All remaining sequence reads were retained as nuclear reads.

The *assemble* and *postprocess* features of Semblans were used separately on the nuclear, mitochondrial, and chloroplast reads, which were assembled independently to generate their respective transcriptomes. The raw reads were first assembled into transcripts using Trinity v2.15.1 (**Grabherr et al., 2011**). Chimeric sequences were then detected from the assembled transcripts using the similarity-based approach developed by **Yang and Smith (2013)** with the DIAMOND v2.1.7.161 algorithm (**Buchfink et al., 2015**). The filtered transcripts were then clustered using Corset v1.09 (**Davidson and Oshlack, 2014**) based on the pseudomapping results from Salmon v1.10.0 (**Patro et al., 2017**). The best-supported sequence among the clusters was retained. From the retained sequences, the open reading frames “ORFs” were predicted using Transdecoder (https://github.com/TransDecoder/TransDecoder). Redundancy in the inferred ORFs for the nuclear dataset was reduced using CD-HIT v4.8.1 (**Fu et al., 2012**) with the settings “-c 0.99 -n 10 -r 0”. The transcriptome completeness was evaluated by measuring the number of recovered single-copy orthologs using BUSCO v5.8.2 (**Simão et al., 2015**), with the “-m transcriptome” setting and the eudicotyledons_odb12 database.

### Homology Inference

Sequence similarity-based homology detection was conducted using all-by-all BLASTN v.2.13.0 (**Altschul et al., 1990**) to infer homolog clusters. The homolog clusters were refined using Markov clustering as implemented in mcl v.14-137 (**Dongen, 2000**) with an inflation value of 1.4. Each inferred homolog cluster was aligned using MAFFT v7.490 (**Katoh et al., 2013**) with the settings “--auto --maxiterate 1000”. The resulting alignments were cleaned using the Phyx (**Brown et al., 2017**) program *pxclsq* with a minimum percent column occupancy threshold set to 10%. Homolog trees were then inferred from the cleaned alignments using Maximum Likelihood (ML) as implemented in IQ-Tree v1.6.12 (**Nguyen et al., 2015**) under the General Time Reversible model of evolution with the gamma parameter to account for among site rate variation “GTR+G”. The inferred homolog trees were filtered using the procedure developed by **Yang and Smith 2014**, where tips on the trees were removed if they had a relative cutoff of 0.2 subs/site or an absolute value cutoff of 0.3 subs/site. The in-paralogs and remaining alternative transcripts were removed from the homolog trees by retaining only one tip for clades consisting solely of ORFs from the same transcriptome. Clades with at least 10 remaining tips separated by a branch of 0.2 subs/site were then divided into separate homolog trees. The coding sequences associated with the resulting homolog trees were then extracted, and the same procedure was repeated twice with the same settings to develop the final homolog trees.

### Ortholog trees, species tree, and molecular dating analysis

We extracted 3366 orthologs with at least 20 taxa from the final homolog trees using the Maximum Inclusion approach with a relative cutoff of 0.2 subs/site and an absolute cutoff of 0.3 subs/site. The sequences belonging to each ortholog tree were aligned using Prank v.170427 (**Löytynoja, 2013**) with default settings. The alignments were cleaned for 30% occupancy using *pxclsq*, and ortholog trees were inferred using IQ-Tree with default model selection (**Kalyaanamoorthy et al., 2017**) and 1000 ultrafast bootstraps (**Hoang et al., 2018**). A coalescent-based species tree was inferred using Astral v.5.7.8 (**Zhang et al., 2018**) with the ortholog trees as inputs. The coalescent branch lengths were converted to molecular branch lengths based on the median subs/site values of concordant ortholog branches using the program Branch Estimation Synthesizer (**Walker et al., 2022**). A supermatrix tree was inferred by concatenating the cleaned alignments using the Phyx program *pxcat*. A ML supermatrix tree with 1000 ultrafast bootstrap (UFBOOT) replicates was inferred using IQ-Tree with linked rate variation among partitions and the GTR+G model of evolution.

The gene trees were rooted using the Phyx program *pxrr,* and the gene shopping SortaDate procedure (**Smith et al., 2018**) was used to identify three genes for molecular dating, ranking them based on concordant bipartitions, root-to-tip variance and tree length, respectively.

Bayesian divergence time inference was conducted in BEAST2 v2.7.7 (**Bouckaert et al., 2019**). Firstly, we used penalised likelihood, implemented in the R package ape (**Paradis et al., 2004**) to convert species tree topology to a starting tree consistent with node constraints. Following **Fishbein et al.** (**2018**), we applied a calibration to the stem lineage of the APSA clade based on a fossil seed bearing a coma assigned to *Apocynospermum*, dated at 47-48 Myr (**Collinson et al., 2012**; **Mertz and Renne, 2005**). We applied a log-normal calibration with an offset of 45 Myr, a mean of 1 and a scale of 1.2, such that 95% of the density lay between 45 and 73 Myr. We further applied a secondary calibration to the root of our tree, based on their results, of a normal distribution with a mean of 54 Myr and a variance of 2.6, such that 95% of the density lay between 49 and 60 Myr. We used a separate GTR+F+G model for each of the three loci, and a separate Optimised Relaxed Clock (**Douglas et al., 2021**) implemented in ORC v1.2.0 for each locus. We kept trees linked and disabled topology proposals to fix the tree. All other priors were left as default. We ran MCMC for 200 million generations, storing trees every 5,000, and verified that all numerical parameters had effective sample sizes > 200 at the end of the run. We used TreeAnnotator to summarise node ages as median heights after burning in 10% of samples.

### Organelle Phylogeny Inference

The predicted ORFs for each organelle were queried against the *Hoya lithophytica* genomes (chloroplast: MW719058.1; mitochondrial: MW719051.1) using BLAST, and the best hit, based upon the length of the match, was retained to ensure each gene for each organelle was only represented by a single ORF. The respective organellar loci were aligned using PRANK with default settings, cleaned using *pxclsq* with a 10% column occupancy minimum, and the cleaned alignments were then concatenated using *pxcat*. Organelle phylogenies were inferred using IQ-Tree with 1000 ultrafast bootstraps and linked rate variation among partitions.

### Gene tree conflict analysis

The coding sequences corresponding to the final homolog trees were aligned using MAFFT with the settings “--auto --maxiterate 1000”, and the resulting alignments were cleaned for 10% minimum column occupancy using the Phyx program *pxclsq*. A ML homolog tree was inferred using IQ-Tree for each cleaned alignment with the GTR+G model of evolution and 1000 UFBOOT replicates for support. The root branch placement for each homolog tree was inferred using the program RootDigger v1.7.0 (**Bettisworth and Stamatakis, 2021**) with default settings. Bipartition conflict between the species tree and the homolog trees was inferred for each node using the program Conflict Analysis and Duplication Inference (CAnDI) (**Robertson et al., 2023**), with the support cutoff set to 95%. CAnDI was also run with no support cutoff to determine the number of times each species tree bipartition had an intersecting bipartition in the homolog trees. Pie charts were produced to visualize the results using the Pies.py script from CAnDI.

### Inference of whole genome duplications and gene duplication

A more stringent redundancy reduction was performed on the inferred ORFs of the nuclear dataset using CD-HIT with the settings “-c 0.90 -n 10 -r 0”. The *dmd* and *ksd* programs from the wgd v2.0.38 (**Chen et al., 2024**) pipeline were used to infer and plot Ks values for each transcriptome.

The extract_clades.py program from **Yang et al. (2015)** was used to extract rooted homolog trees from the unrooted conflict analysis homolog trees that met the criteria of duplicating at the root of the ingroups. Gene duplications occurring at each node of the species tree were inferred using the program PhyParts v0.0.1 (**Smith et al., 2015**) with verbose mode. The rooted homolog trees were annotated by extracting the single longest amino acid sequence corresponding to a tip in each tree. We then conducted a BLASTP analysis of the sequences against a database of *Arabidopsis thaliana* (TAIR10) peptides to identify the best hit with a minimum e-value of 1e-3. Genes without a hit to *A. thaliana* were run against the National Center for Biotechnology Information Non-Redundant database on 04-13-2025.

For five nodes of interest, we used the Arabidopsis IDs associated with each gene family that expanded at the node to perform a gene ontology (GO) analysis. The analysis was conducted using PANTHER19 (**Thomas et al. 2022**) with the GO database (10.5281/zenodo.14861039 Released 2025-02-06). The analysis was performed using the overrepresentation test (**Mi et al., 2019**) (Released 20240807) and the “GO biological process complete” dataset. Fisher’s exact test was performed with a Bonferroni correction for multiple testing.

To further investigate the gene duplication history of *CYP87A*, we translated the coding sequences from the *CYP87A* homolog “cluster11016_1rr_1rr_1rr_1rr” into peptides, aligned them using PRANK, and cleaned the alignments with pxclsq for a minimum column occupancy of 30%. We inferred the model of evolution and constructed a ML tree using IQ-Tree. We then performed the same process on a dataset that contained both our homolog data and that used by **Feng et al. (2025)**. Their dataset was composed of homologs of the *CYP87A* gene family which was used to document the gene family’s evolution in *Asclepias curassavica*. In addition to *A. curassavica*, their data included *CYP87A* genes from *Asclepias syriaca*, *Calotropis gigantea*, *Calotropis roseus*, *Erysimum cheiranthoides*, *Digitalis lanata*, *Nicotiana benthamiana*, *Solanum lycopersicum*, *Arabidopsis thaliana,* and *Oryza sativa*.

## Results

### Dataset Composition

The dataset included 48 ingroup species encompassing four of the five Apocynaceae subfamilies, excluding Secamonoideae. Sampling captured 13 tribes and 28 genera. The number of raw sequence reads ranged from 1,165,862 for *Hoya macgillivrayi* to 180,045,873 for *Catharanthus ovalis* (**Supplementary Table 1**). After processing and quality filtration of the reads, the lowest level of detected contamination was 2.28% for *Rauvolfia serpentina,* and the highest was 50.3% for *Pachypodium lamerei*. The proportion of reads identified as nuclear sequences was between 20% for *Asclepias viridiflora* and 94% for *Catharanthus roseus*. The proportion of chloroplast reads ranged from 0.01% for *Lacmellea panamensis* to 17.5% for *A. viridiflora,* while the mitochondrial reads ranged from 0.008% for *Calotropis gigantea* to 1.56% for *H. macgillivrayi*.

The number of predicted nuclear Open Reading Frames (ORFs) in the resulting assemblies ranged from 4148 for *H. macgillivrayi* to 79,190 for *Plumeria rubra*. The number of genes predicted for the chloroplast ranged from 13 for *Lacmellea panamensis* to 41 for *Cynanchum japonicum*, *A. viridiflora* and *Rhazya stricta*. No mitochondrial genes were assembled for *Spigelia splendens*, *Vinca major*, *Asclepias curassavica* and *Hoodia gordonii*, and 30 genes, the most of any species, were assembled for *Wrightia religiosa.* Of the 2805 single-copy orthologs in the eudicotyledons BUSCO database, as few as 52 were present for the *H. macgillivrayi* transcriptome and as many as 2549 for the *Catharanthus ovalis* transcriptome (**Supplementary Figure 1**).

### Inferred Species Relationships and Divergence Dates

Complete concordance with perfect support for all species relationships was inferred between the concatenated maximum likelihood and coalescent-based trees (**Supplementary Figure 2**). All tribes were recovered as monophyletic, and all genera formed a clade aside from *Ceropegia* (**Figure 1**). The *Ceropegia* and *Hoodia* formed a clade subtending a shift in molecular rate, where the branch is 0.0688 subs/site, and the sister to the branch is 0.0069 subs/site (**Figure 2A**). This shift in molecular rate affected most of the concordant genes (**Figure 2B**).

**Figure 1.**
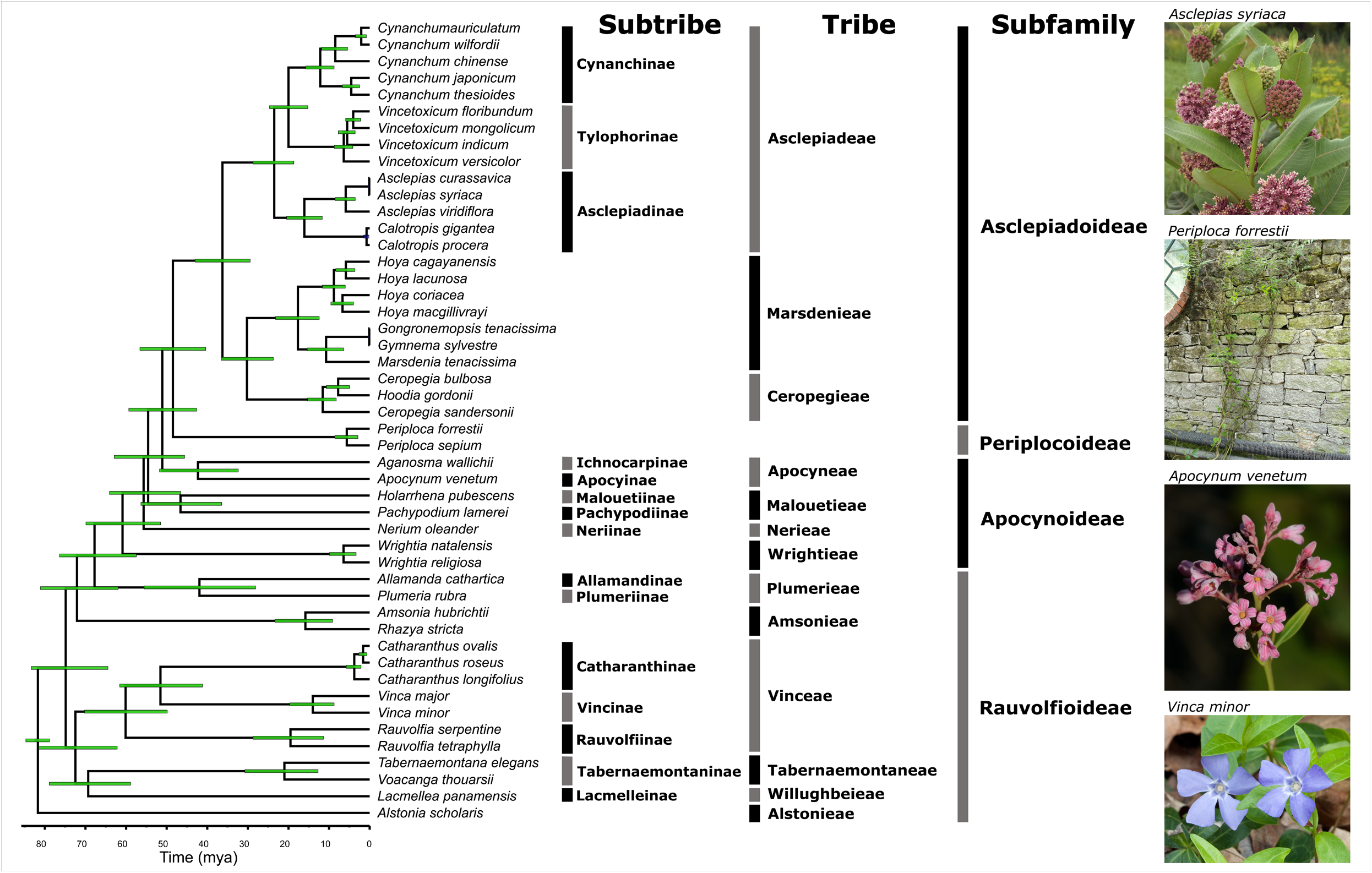
The inferred relationships in the Apocynaceae with the dates were applied using the gene shopping Bayesian dating analysis. The bars overlaying the divergences represent the 95% HPD intervals. *Photo credits*: *Periploca forrestii*; Daderot, *Apocynum venetum*; Gideon Pisanty, *Asclepias syriaca*; Amos Oliver Doyle, *Vinca minor*; Ryan Kaldari. Licenses and location of original photographs can be found in **Supplementary Table 3**.

**Figure 2.**
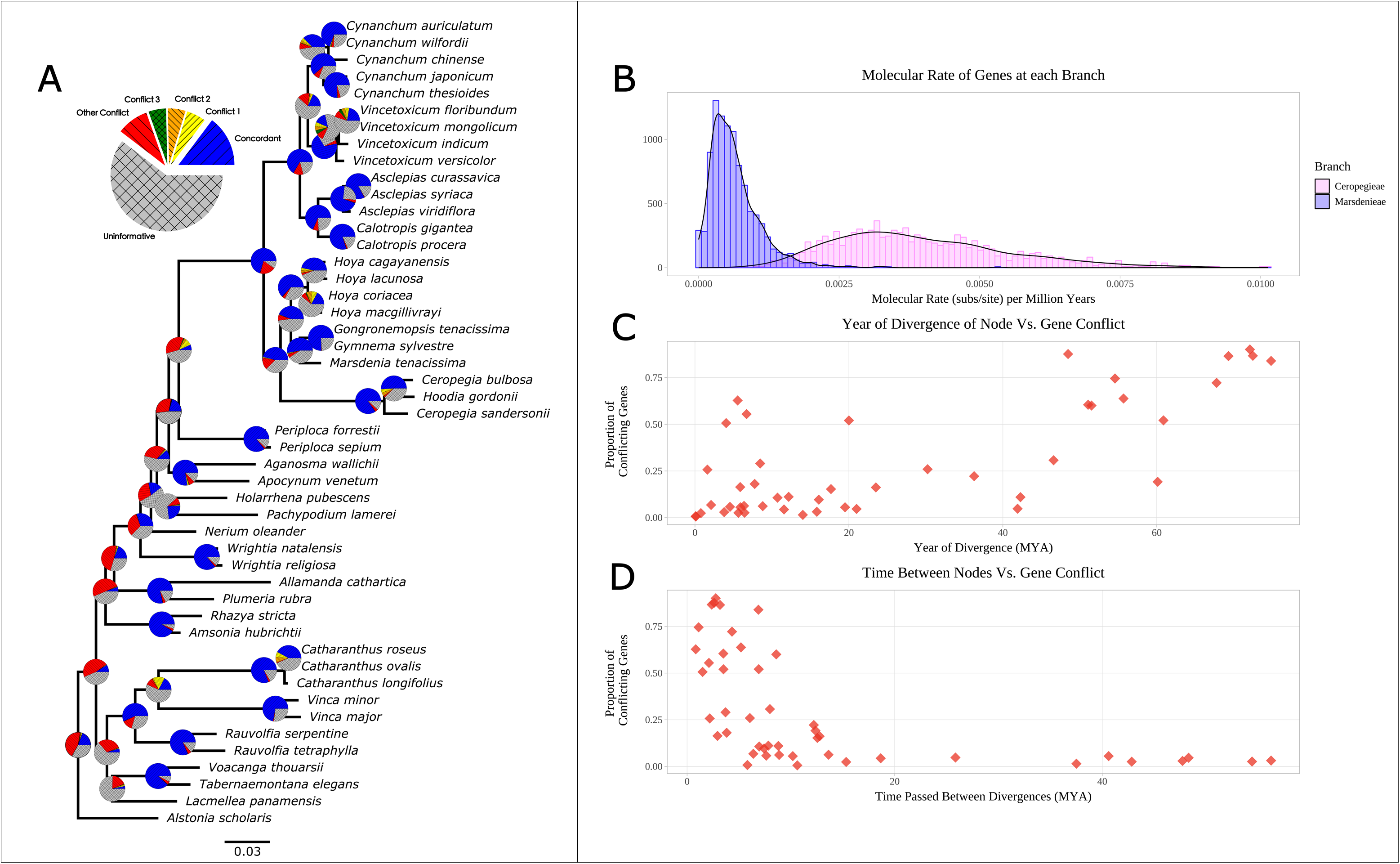
Inferred gene tree conflict and rate variation in the Apocynaceae. **A)** Phylogram of the inferred species relationships with branches in subs/site. Pie charts depict the proportion of gene trees inferring a relationship, with blue being concordant, yellow, orange, green being the first, second, and third most common conflicts, and all other conflicts in red. All relationships inferred to be uninformative, lacking necessary sampling or UFBoot <= 95, are in grey. **B)** Density plot of the distribution of molecular branch lengths for concordant ortholog trees converted to subs/site/million years, with purple being the branch leading to Marsdenieae and pink being the branch leading to Ceropegieae. **C)** The relationship between the time between divergences and the gene trees with supported conflict. **D)** The relationship between the divergence time and the gene trees with supported conflict.

The subfamily Rauvolfioideae was inferred to be paraphyletic. Within the subfamily, the tribe Alstonieae was recovered as sister to all other Apocynaceae. The tribes Willughbeieae, Tabernaemontaneae, and Vinceae form a monophyletic group, with Tabernaemontaneae and Willughbeieae forming a clade sister to tribe Vinceae. Within Vinceae, the subtribe Rauvolfiinae is sister to a clade containing the subtribes Catharanthinae and Vincinae. Paraphyly of Rauvolfioideae was also caused by Amsonieae and Plumerieae forming a grade leading into the Apocynoideae.

The Apocynoideae was inferred to have diverged ∼60 mya and recovered to be paraphyletic, consisting of a grade of the sampled tribes Wrightieae, Nerieae, Malouetieae, and Apocyneae diverging in that order and leading to a clade with the subfamilies, Periplocoideae and Asclepiadoideae as sister to one another, diverging ∼48 mya. The Asclepiadoideae diverged ∼36 mya, and within the clade, the tribes Marsdenieae and Ceropegieae form a clade sister to Asclepiadeae. The monophyletic Asclepiadeae contains Asclepiadinae sister to a clade of Cynanchinae and Tylophorinae.

### Gene tree and organelle conflict

The mitochondrial phylogeny showed low ultrafast bootstrap support (UFBOOT < 95) for, and conflicted with, almost all relationships recovered by the nuclear phylogeny (**Supplementary Figure 3**). The relationships inferred from the chloroplast data were predominantly highly supported (95 <= UFBOOT), except for the Rauvolfioideae intersubfamilial relationships (**Supplementary Figure 3**). The chloroplast tree was highly congruent, with a few notable exceptions. In the chloroplast tree, Rauvolfiinae was recovered as sister to Vincinae with low support. This relationship was the only common alternative conflict among the nuclear gene trees supported by 1872 gene trees in comparison to the nuclear relationship that was supported by 2247 gene trees (**Figure 2A**).

The chloroplast tree also recovered Cynanchinae as sister to a clade of Asclepiadinae and Tylophorinae, and this relationship did not have a common conflict among the nuclear gene trees. The Marsdenieae tribe (Asclepiadoideae) was inferred to be paraphyletic. *Hoya lacunosa* and *Marsdenia tenacissima* were well supported to be separated from the rest of the Marsdenieae tribe, forming a clade with the Ceropegieae tribe. This relationship places the Ceropegieae tribe as nested within Marsdenieae.

Five relationships had two alternate conflicting topologies of nearly equal proportion, suggesting that ILS was the predominant source of conflict. Of note were the intergeneric relationships in *Catharanthus, Hoya, Vincetoxicum, and Cynanchum*. The sister relationship of *Hoodia gordonii* and *Ceropegia bulbosa* also demonstrated this pattern of conflict. The backbone of the Apocynaceae showed the highest level of conflict without dominant alternatives, with all relationships showing close to equal or greater levels of total conflict compared to the genes showing concordant relationships (**Figure 2A**).

Among the nuclear gene trees, the backbone relationships of the phylogeny had the highest proportion of conflicting gene tree signals. Increased supported conflict was observed for the divergences in the backbone (**Figure 2C**). We recovered a general pattern that the shorter the time between divergences, the greater the proportion of conflicting gene trees (**Figure 2D**).

### Gene family expansion and genome duplication

In total, 14,838 gene families containing at least four different taxa were inferred to have expanded at the base of Apocynaceae. Within the family, the clade comprising *Gongronemopsis tenacissima* and *Gymnema sylvestre* had the greatest number of shared duplications, with 4,967, and the genus *Vinca* had the second greatest number, with 4,154 (**Figure 3A**). Between 11 and 430 duplications were inferred at the divergences along the backbone, and the Asclepiadoideae together shared 1,625. The genome duplication history remains difficult to interpret through Ks plots and gene duplications alone, but both *Vinca* species exhibited a similar peak at around 0.25 Ks (**Figure 3C; Supplementary Figure 4**).

**Figure 3.**
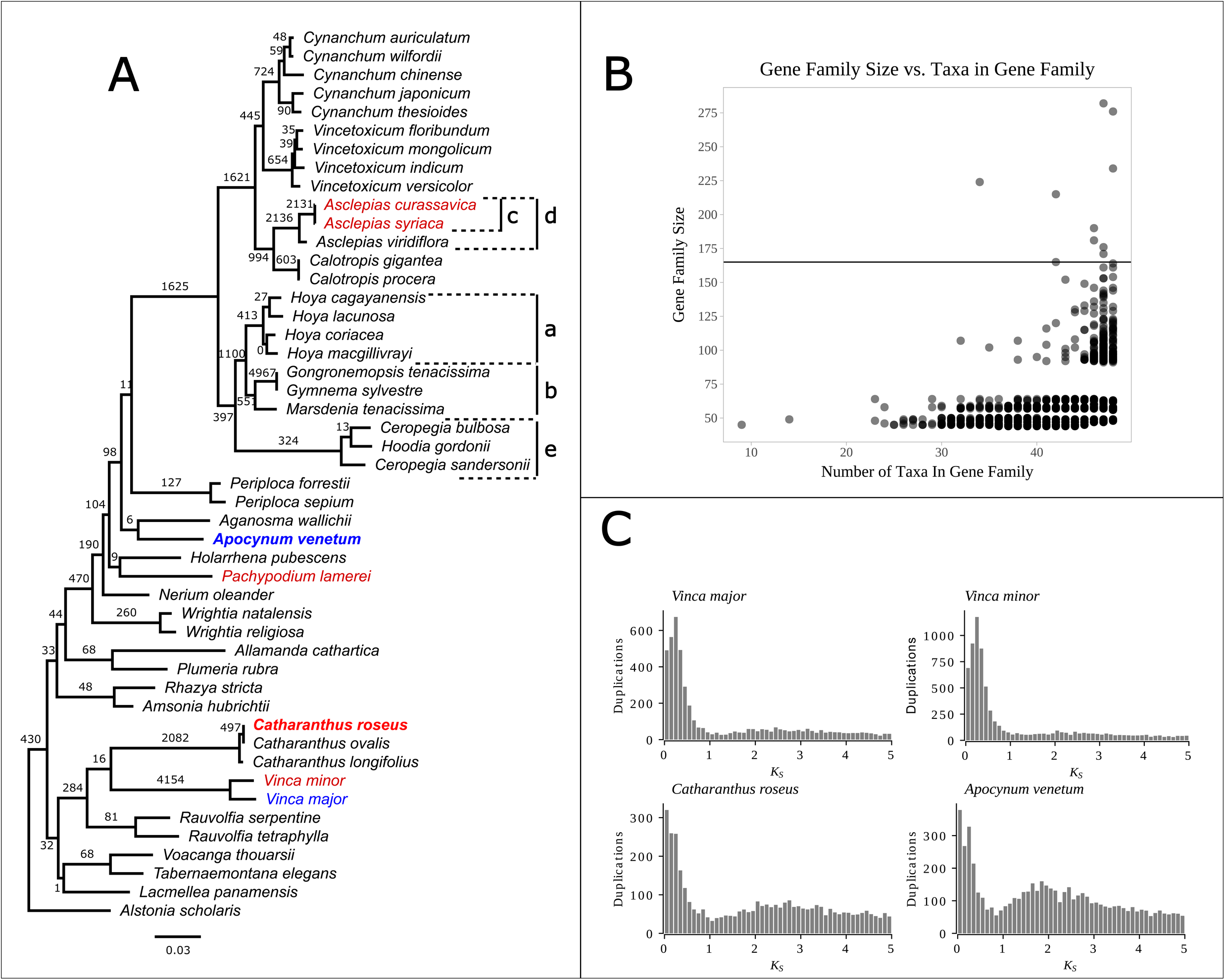
Gene and genome duplications in the Apocynaceae. **A)** Inferred phylogram with number of inferred gene duplications labeled on each branch. Taxa in red are those with a paper reporting that they have not experienced a whole genome duplication or a whole genome duplication was not investigated, taxa in blue are those predicted to have experienced a genome duplication, and bolded taxa are those inferred to share a genome duplication. **B)** The relationship between the size of a gene family that expanded at the base of Apocynaceae and the number of taxa with that gene family. The ten largest gene families are above the line. **C)** Ks Plots for the species discussed in the text.

The ten largest gene families that expanded at the base of the Apocynaceae ranged from 165 tips, with representatives from 42 different taxa, to 282 tips, with representatives from 47 different taxa (**Figure 2B**; **Table 1**). In general, the larger gene families contained more representative taxa except for the “*S-locus lectin protein kinase family protein”*, which had 224 gene copies across only 34 representative taxa.

**Table 1.**
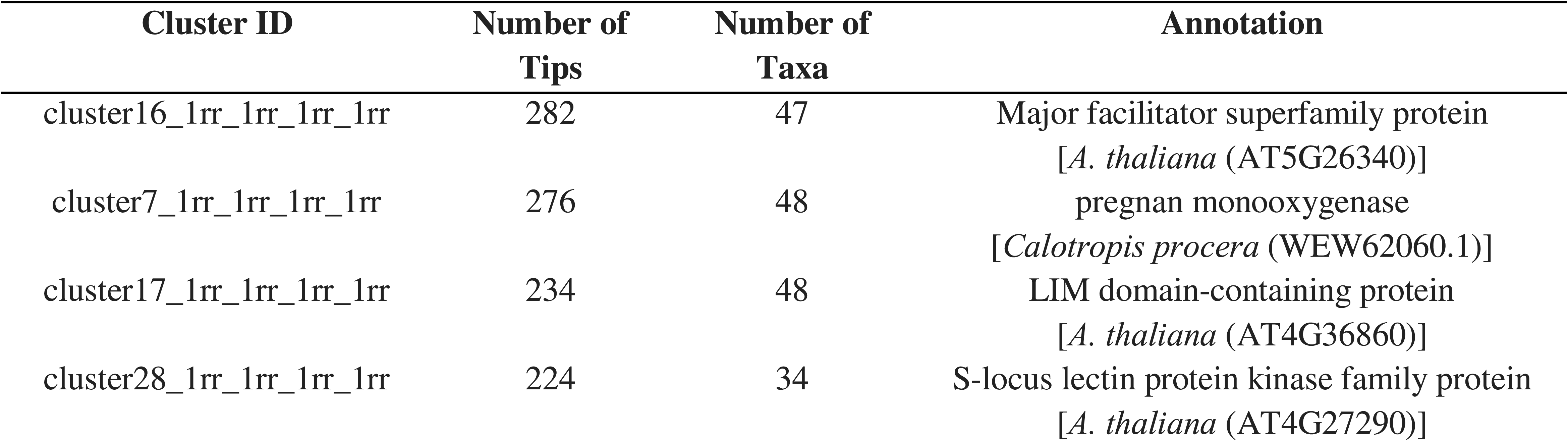
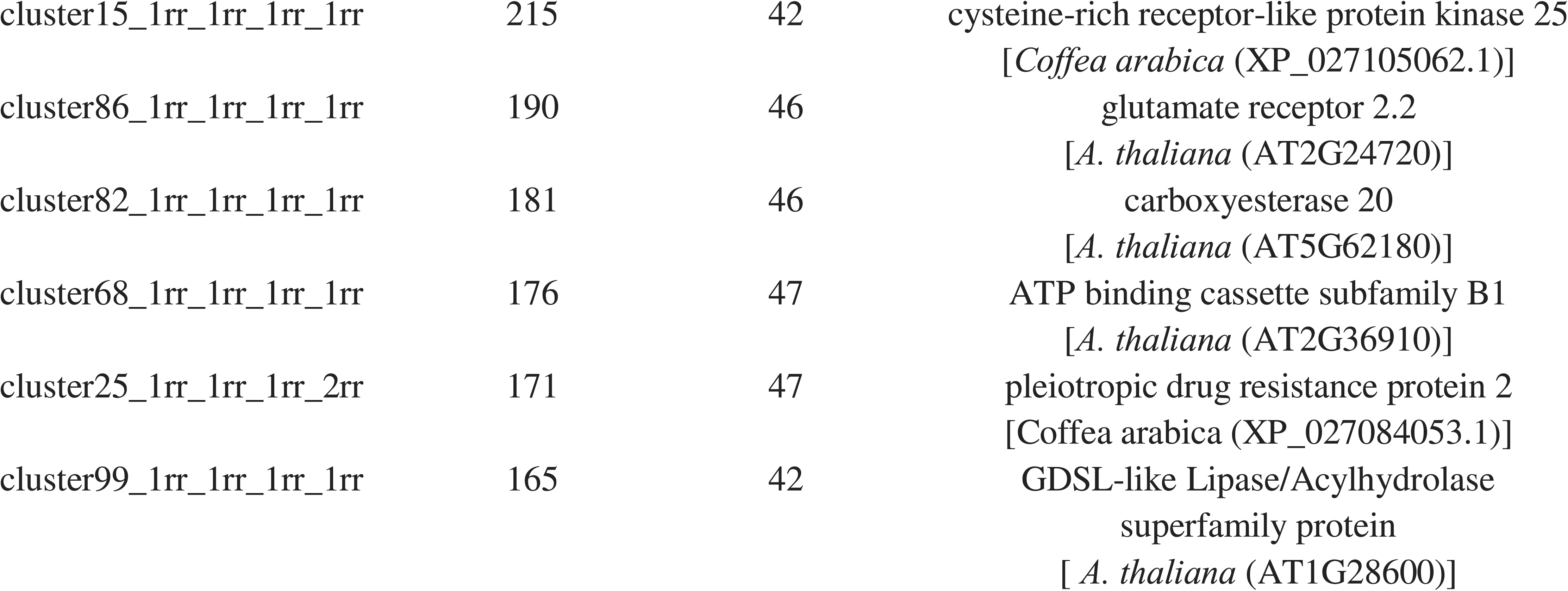
The ten largest gene families that have expanded in the Apocynaceae.

| Cluster ID | Number of Tips | Number of Taxa | Annotation |
| --- | --- | --- | --- |
| cluster16_1rr_1rr_1rr_1rr | 282 | 47 | Major facilitator superfamily protein<br>[*A. thaliana* (AT5G26340)] |
| cluster7_1rr_1rr_1rr_1rr | 276 | 48 | pregnan monooxygenase<br>[*Calotropis procera* (WEW62060.1)] |
| cluster17_1rr_1rr_1rr_1rr | 234 | 48 | LIM domain-containing protein<br>[*A. thaliana* (AT4G36860)] |
| cluster28_1rr_1rr_1rr_1rr | 224 | 34 | S-locus lectin protein kinase family protein<br>[*A. thaliana* (AT4G27290)] |
| cluster15_1rr_1rr_1rr_1rr | 215 | 42 | cysteine-rich receptor-like protein kinase 25<br>[*Coffea arabica* (XP_027105062.1)] |
| cluster86_1rr_1rr_1rr_1rr | 190 | 46 | glutamate receptor 2.2<br>[*A. thaliana* (AT2G24720)] |
| cluster82_1rr_1rr_1rr_1rr | 181 | 46 | carboxyesterase 20<br>[*A. thaliana* (AT5G62180)] |
| cluster68_1rr_1rr_1rr_1rr | 176 | 47 | ATP binding cassette subfamily B1<br>[*A. thaliana* (AT2G36910)] |
| cluster25_1rr_1rr_1rr_2rr | 171 | 47 | pleiotropic drug resistance protein 2<br>[*Coffea arabica* (XP_027084053.1)] |
| cluster99_1rr_1rr_1rr_1rr | 165 | 42 | GDSL-like Lipase/Acylhydrolase<br>superfamily protein<br>[*A. thaliana* (AT1G28600)] |

A focused analysis of the Gene Ontology terms for five nodes where a gene family expanded at the base of the node showed that genes associated with changes in trichomes were disproportionately duplicated in the epiphytic clade *Hoya* (A). The divergence prior to that (B) did not show similar enrichment, with the greatest fold change affecting genes associated with DNA repair and damage (**Table 2**). The gene duplications for the clade of *Asclepias syriaca* and *A. curassavica* (C) showed a higher rate of duplication for genes associated with immunity, and the node inclusive of *A. viridiflora* (D) did not show the enrichment of genes related to immunity. The genes showing duplications at the base of the *Ceropegia* clade (E) were predominantly associated with metabolic processes.

**Table 2.**
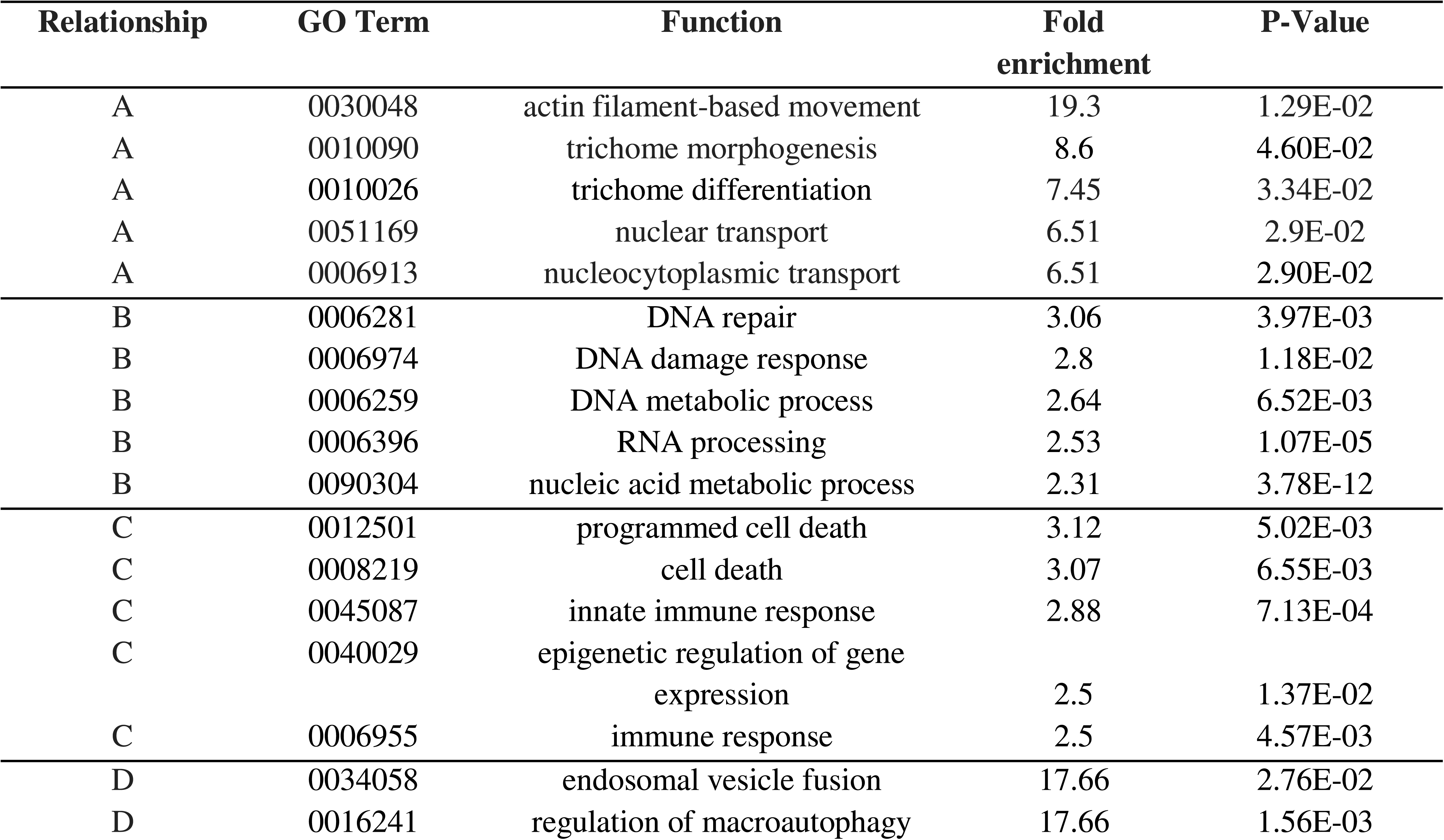
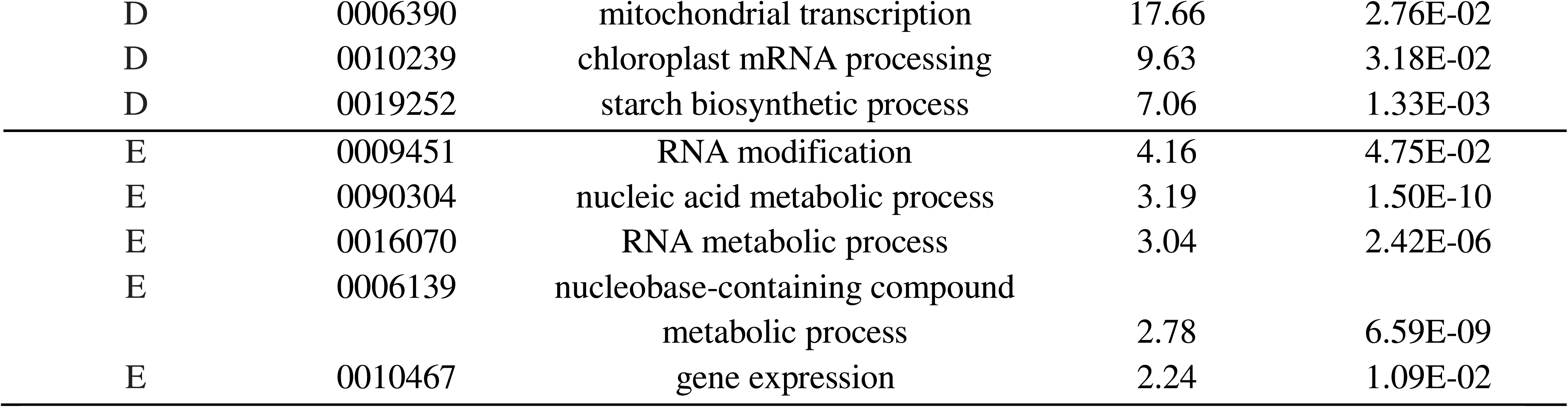
Gene expansion with relation to Gene ontology analyses (Biological Process). The terms with the five highest folder changes are presented. The complete list may be found in Supplementary Table 2.

We further examined the *CYP87A* gene tree, finding that a series of gene duplications were inferred to have occurred at the base of cardenolide producing *A. syriaca* and *A. curassavica* species (**Figure 4A**). Based on our data alone, the duplication happens after the split with *A. viridiflora*. However, after combining our data with the data from **Feng et al., 2025**, we found *A. syriaca* and *A. curassavica* shared the same ancestral gene copy found in *A. viridiflora* (**Figure 4B**). However, we could not recover any additional copies of *CYP87A* from *A. viridiflora* when comparing the transcriptome sequences to the protein sequences with BLAST. This indicates the additional *CYP87A* copies were either lost, or not expressed at the time of sampling.

**Figure 4.**
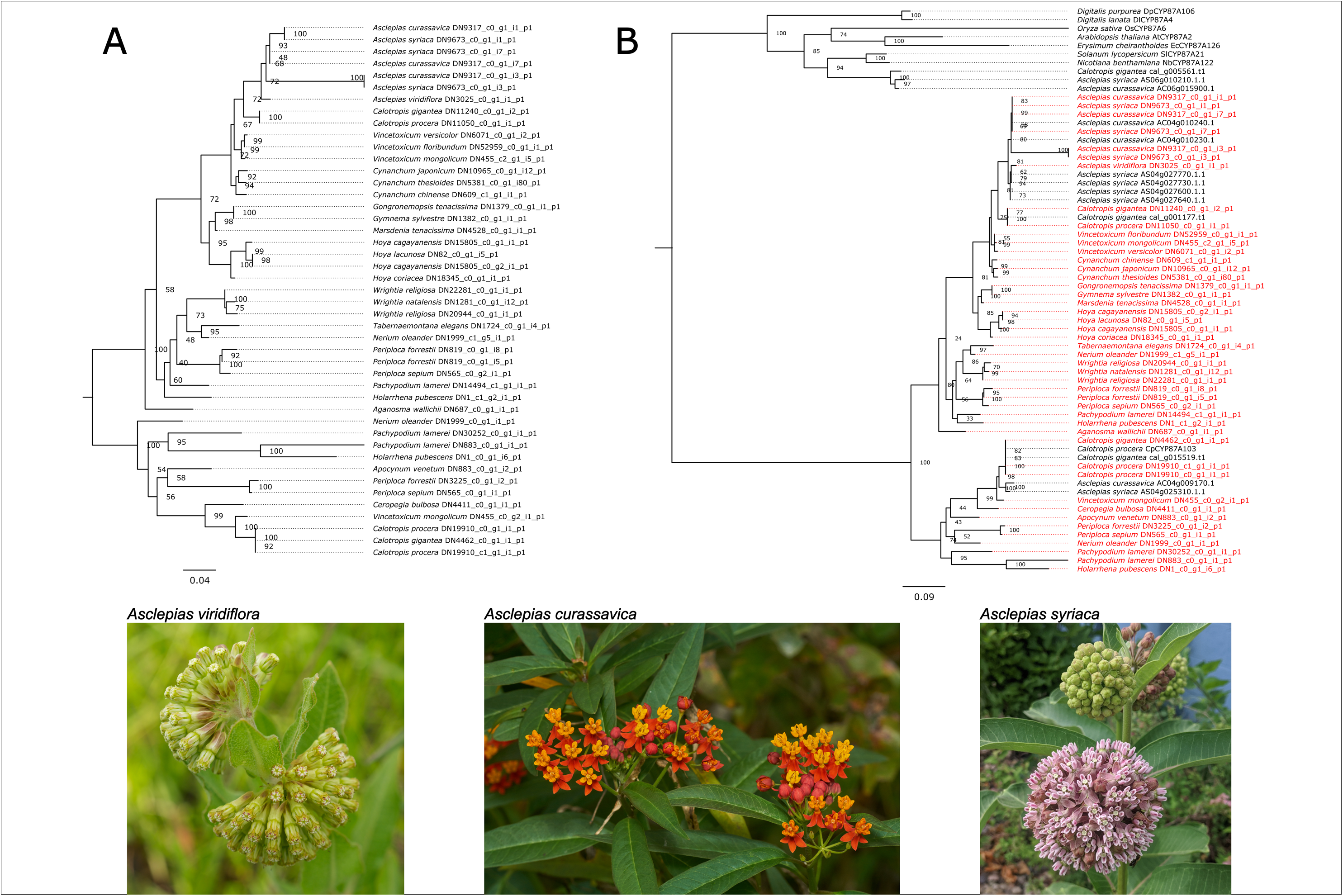
Evolutionary history of *CYP87A* in the Apocynaceae. **A)** The gene tree of CYP87A using only data from this study. **B)** The gene tree of *CYP87A* with the data in this study (colored in red) and the data from Feng et al., 2025 (in black).

## Discussion

### The use of publicly available transcriptomes for phylotranscriptomics

More than 6000 plant species have publicly available transcriptome sequence data. As noted by **Wong and Peakall, 2022**, transcriptome data from differential gene expression studies is an excellent source of data for phylotranscriptomics. The dogbane family, Apocynaceae, produces diverse biochemical compounds, and the species studied are phylogenetically spread across the clade and have been the focus of many differential gene expression studies. The dataset used in this paper consisted of 55 recycled transcriptomes, predominantly sourced from differential gene expression studies, along with one newly sequenced transcriptome obtained from silica-dried tissue.

As expected, given the data was sourced from a wide variety of studies conducted through the years and across the globe, the number of inferred ORFs and BUSCOs varied greatly among samples. Aside from *Hoya macgillivrayi*, which had the fewest initial reads, all samples had at least 50% of BUSCOs recovered, with some recovering more than 90%. Recycling transcriptome data provides a cost-effective approach to gain preliminary data for future work, real data for student-led projects, and navigating uncertain funding climates. Many of the insights and analyses from this project could have arisen without our newly sequenced *A. viridiflora* transcriptome. However, the successful use of this sample exemplifies how silica-dried (**He et al., 2022**; **Ruiz-Vargas et al., 2024)** or herbarium-pressed tissue (**Tyszka et al., 2024; Tyszka et al., 2025**) provide a source of novel transcriptomes, enhancing studies primarily utilizing publicly available data and alleviating fieldwork costs.

### Phylogenetic relationships of the Apocynaceae

The phylotranscriptomic inference presented here is the most gene-dense examination of Apocynaceae to date, complementing species-rich studies (**Fishbein et al., 2018; Wang et al., 2023**). Notably, the dataset includes nuclear genes, providing novel insights beyond those of the predominantly chloroplast-based earlier research. The chloroplast is often considered to contain a single evolutionary history (**Doyle, 1992**; **Doyle, 2022**), with some inferred exceptions (**Walker et al., 2019**; **Goncalves et al., 2019**). Similarly, the mitochondrial genome is typically thought of as having a single point of coalescence and is theoretically, though not always in practice, shared with the chloroplast (**Tyszka et al., 2023; Liang et al., 2025; Dominicus et al., 2025**). However, the sparse gene recovery likely introduced misleading phylogenetic signals in this study.

Species tree inference using the nuclear genes revealed complete concordance between the coalescent-based and supermatrix topologies. Such concordance is uncommon in phylotranscriptomic studies, as the two inference methods treat gene evolutionary histories in divergent ways. The coalescent-based method uses quartets to properly infer relationships in the anomaly zone, where the most common gene tree relationship does not match the species tree due to ILS (**Degnan and Rosenberg, 2006**). Furthermore, while the supermatrix approach provides stronger signal for relationships, it can give genes disproportionate influence over the inferred topology (**Gatesy and Baker, 2005**; **Shen et al., 2017**), a problem exacerbated by misidentified orthology (**Brown and Thompson, 2017; Walker et al., 2018)**.

The topology recovered in this study is similar to recent Apocynaceae systematic studies and supports the paraphyly of Rauvolfioideae and Apocynoideae (**Sennblad and Bremer, 1996**; **Rapini et al., 2007**; **Fishbein et al., 2018**; **Antonelli et al., 2021**). However, the placement of Periplocoideae conflicts with several previous investigations (**Supplementary Figure 5**). Historically, this subfamily has been recovered as nested within (**Sennblad and Bremer, 1996**; **Fishbein et al., 2018**; **Antonelli et al., 2021**) or diverging before Apocynoideae (**Rapini et al., 2007**). We inferred the divergence of Periplocoideae as after Apocynoideae, placing it sister to Asclepiadoideae. However, because our dataset lacks several groups within Apocynoideae (e.g., tribe Baissieae), this placement does not directly conflict with all previous studies. Furthermore, high levels of gene tree conflict at the Periplocoideae-Apocynoideae node show that underlying discordance may explain the contention.

One notable difference between the chloroplast and nuclear topology is a molecular rate shift on the branch leading to a clade of *Ceropegia* and *Hoodia*. In plant phylogenetics, molecular rate shifts in phylogenies often reflect life history shifts (**Smith and Donoghue, 2008**; **Yang et al., 2015**). However, this is not the case based on the group’s extant taxa. The shift in molecular rate leading to *Ceropegia* and *Hoodia* seems predominately correlated to morphological change, specifically as *Hoodia* is cactus-like. The low levels of gene tree conflict for the monophyly of *Ceropegia* and *Hoodia* would indicate that a large portion of the genome was able to fix, and potentially, this group underwent a decrease in population size. The rate shift was not limited to specific genes, as most genes concordant with the relationship show a rate shift in that particular branch **(Figure 2B)**. Although difficult to tell from the three taxa in this dataset alone, the potential underlying reason for the *Ceropegia/Hoodia* rate shift is not easily explained and is worth examining in future studies.

The Asclepiadeae-Asclepiadoidea clade shows conflict between the chloroplast and nuclear inferred topologies. The topology inferred by the chloroplast data is supported by the findings of **Fishbein et al. (2018)**, who used plastid genes and whole plastome data to infer their species tree. This suggests that cytonuclear discordance in Apocynaceae is likely a biological signal (**Rieseberg et al., 1991**; **Larson et al., 2024**). Although backbone concordance is found between studies, gene tree conflict is especially prevalent along these nodes, and indicates that complex dynamics, such as introgression and ILS, were at play during the speciation process.

### Relationship of gene tree conflict and time

Gene tree conflict provides a rare connection between micro- and macroevolutionary events, where a change that once occurred in an individual may be inferred from genomic data millions of years later. Processes that generate conflict include hybridization, gene duplication and loss, incomplete lineage sorting, and horizontal gene transfer. Gene tree conflict has been correlated with rapid radiations and morphological innovations (**Parins-Fukuchi et al., 2021**; **Stull et al., 2021**), making it especially prominent during periods of major evolutionary change.

The greatest proportion of gene trees exhibiting conflicting signals was found in the older divergences that comprise the backbone of the Apocynaceae, previously noted for its rapid radiation (**Straub et al., 2014**). These divergences were inferred to have occurred over short time spans. Since increased conflict can be a signal of rapid radiation due to a lack of gene fixation prior to speciation, we tested whether the time between divergences correlated with the proportion of conflict. We found that the less time predicted to have passed, the greater the proportion of genes that show supported conflict. This may indicate that the backbone relationships of Apocynaceae historically had a genetically diverse and large population size, leading to successive speciation events.

### Gene duplication and gene family expansion in the Apocynaceae

Phylotranscriptomic datasets are a cost-effective approach for identifying genes of interest. These datasets can be used to infer the expansion of a gene family within a clade, an important source of novel genetic material upon which selection can act. The gene families inferred to have expanded at the base of Apocynaceae typically show a correlation between the size of the family (as measured by the number of tips) and the number of taxa (**Figure 3B**; **Table 1**). The exception is the “*S-locus lectin protein kinase family protein*” involved in self-incompatibility and known for complex expression patterns (Ramanauskas and Igić**, 2021**; **Ramanauskas et al., 2025**). The gene family was not recovered in most species of Rauvolfioideae, aside from the clade containing *Voacanga thouarsii* and *Tabernaemontana elegans* (**Supplementary Figure 6**). The gene was also not recovered in *Pachypodium lamerei.* We cannot reject that the S-locus gene family was not recovered in the Rauvolfioideae family due to potential methodological issues arising from the rapid rate of evolution in the gene family or potential lack of expression in the tissue sampled. However, it may result from a complex pattern of S-locus evolution and warrants investigation with genomic data.

The branch leading to the genera *Ceropegia* and *Hoodia* exhibited a shift in molecular rate, which correlates with their comparatively extreme morphological changes. We examined the Gene Ontology (GO) terms for gene family expansions specific to this clade to identify enrichment for terms associated with morphological change. No terms associated with morphological features were present in the five GO terms experiencing the greatest fold change. However, our sampling only included one *Hoodia*, whose transition to a cactus-like form is a more pronounced morphological change than *Ceropegia*.

We also investigated the transition to epiphytism, which occurred at the base of the *Hoya* genus. Notably, among GO terms with the highest fold change were those associated with trichome morphogenesis, a trait that has been the focus of recent work in the *Hoya* (**Basir et al., 2022**). Furthermore, trichomes have been shown to play an important role in the transition to epiphytism (**Ha et al., 2021**), and across a broad range of families, this life history shift has been associated with trichome modifications for foliar absorption (**Li et al., 2023**). Overall, the alterations to trichomes are expected to change the life history to epiphytism, and in the *Hoya* genus, gene family expansion may have facilitated this transition.

The milkweed species *Asclepias syriaca* and *Asclepias curassavica* are both known to produce cardenolides as a defense mechanism against herbivory. In contrast, one report demonstrated that *Asclepias viridiflora* does not produce cardenolides (**Keeler and Tu, 1983**). Cardenolide production has been attributed to protection against herbivory. Therefore, not investing in their production is correlated with lower rates of predation (**Pellissier et al., 2014**). When examining the GO terms associated with duplications specific to the *A. syriaca* and *A. curassavica* clade, four of the top five with the highest fold change were associated with either immunity or cell death. The gene expansions occurring specific to all *Asclepias* species did not show this same pattern, warranting further investigation into the role of gene duplication in cardenolide production.

Cardenolides are among the most studied defensive compounds produced in *Asclepias* in part due to their medicinal properties. Part of the biosynthetic pathway involves pregnane derivatives (**Hassan et al., 2023**; **Kunert et al., 2023**), one of the most expanded families in the Apocynaceae (**Table 1**). The *CYP87A* gene has recently been a focus of studies into cardenolide production inside (**Feng et al., 2025**) and outside (**Kunert et al., 2023**) the Apocynaceae. The gene tree for the *CYP87A* gene was previously shown to have *Asclepias*-specific duplications (**Feng et al., 2025**). Here, we examined whether that same duplication was in the non-cardenolide producing *A. viridiflora*. In the tree consisting solely of our data, we found that *A. viridiflora* contained a single copy of *CYP87A*, which was sister to the duplicated copies found with *A. syriaca* and *A. curassavica*. Incorporation of the **Feng et al., 2025** dataset found that the *A. viridiflora* copy of *CYP87A* was present in *A. syriaca* and *A. curassavica*. When examining the transcriptome of *A. viridiflora* for evidence of the additional copies of *CYP87A,* we still only recovered a single copy. Despite extraction from silica dried tissue, the *A. viridiflora* transcriptome had the best BUSCO scores, indicating a slightly higher level of completeness. Although we could not recover copies of *CYP87A* from *A. viridiflora*, they may still be present in the genome, and future genomic work should investigate the complex association between *CYP87A* duplication and cardenolide production in the *Asclepias* genus.

### Whole genome duplications in Apocynaceae

Inferring whole genome duplications is a complex and potentially fallible process, as such, this could explain recent reports in genomic work (**Tasdighian et al., 2025)**. One notable genome duplication that has been inferred is in *Vinca major* (**Obermayer and Greilhuber, 2005**; **Ren et al., 2018**). However, the closely related *Vinca minor* is reported as being diploid in the genome study (**Stander et al., 2022**). Based upon the discussion of Cryptoploidy within *Vinca* by **Obermayer and Greilhuber, 2005**, evidence points to both of these being correct. Within our data, without the context of these studies, the large number of shared duplications and the similar Ks peaks (**Figure 3**) would be evidence that a shared genome duplication has occurred. This is an indication that by Ks plot and gene duplication alone, it is easy to be misled (**Kellogg, 2016; Tasdighian et al., 2025**), and as more lines of evidence become available hypotheses about whole genome duplications will likely be updated.

As genome assemblies provide one of the most reliable means of testing genome duplication (**Kellogg, 2016**), they are often taken as definitive. The genome study of *Apocynum venetum* (**Dorjee et al., 2024**) showed evidence for a shared duplication with *Catharanthus roseus*. As presented in our inferred Ks plot, a small peak exists in *A. venetum* data. However, the same Ks peak does not appear in *C. roseus* (**Figure 3C**). **Xu et al. (2023)** also find little evidence supporting a whole genome duplication in *C. roseus.* The shared peak reported by **Dorjee et al., 2024**, would indicate a WGD at the base of Apocynaceae based on the phylogenetic structure. If this were to be the case, we would expect to see evidence of a WGD among all taxa that share this node. Our Ks plots show little evidence for WGDs among these taxa. Additionally, genome studies of *Asclepias curassavica* (**Feng et al., 2025**), *Asclepias syriaca* (**Weitemier et al., 2019**), *Pachypodium lamerei* (**Cuello et al., 2024**) and *Voacanga thourasii* (**Cuello et al., 2022**) either do not discuss a WGD or did not find evidence for one in the data indicating that a shared WGD may not be supported. As mentioned, WGD inference by Ks plot and gene duplication alone may be misleading (**Kellogg, 2016; Tasdighian et al., 2025**). The contradiction among lines of evidence suggests that as additional genome assemblies become available, it will be important to determine the existence of a whole genome duplication along the branch succeeding the divergence of Alstonieae. Further debates on the position and existence of genome duplications will likely continue into the era of WGD detection aided by synteny information.

## Conclusions

This study highlights the value of recycling transcriptome data to assemble phylotranscriptomic datasets for insights into molecular evolution. We used predominantly publicly available data to conduct the first phylotranscriptomic analysis of the dogbane family. The transcriptome data revealed the complexity and challenges of inferring the group’s whole genome duplication history. We also uncovered evidence that gene duplication may have facilitated evolutionary adaptations within the clade. The nuclear data revealed pervasive gene tree conflict along the backbone relationships of the family, and we identified a correlation between divergence time intervals and the amount of conflict. We encourage researchers to leverage publicly available data to investigate clades that have not yet been the focus of phylotranscriptomic work.

## Author Contributions

S.A, J.F.W and J.H-M led the experimental design with input from all authors. J.F.W and S.A performed the transcriptome assembly and data accumulation. NWH performed the molecular dating analysis. Data analysis was led by S.A and J.F.W with assistance from M.R.C, S.B, E.P, M.R, C.S, N.M, A.D, B.D.N, V.M, C.S, A.D, D.A, Z.H, J.N.P, S.D, T.M, T.M, B.M.M.S, N.T, S.D-H, and input from N.W-H and J.H-M. Figure generation was led by S.A, with input from J.F.W and B.D-N. SA and JFW led the writing, with input from all other authors. All authors approved the final draft of the manuscript.

## Acknowledgements

We thank Mary Ashley for providing the *A. viridiflora* tissue and for the helpful discussion regarding the species. Alexa Tyszka for doing the RNA extraction of *A. viridiflora*. We thank Zarema Arbieva and Nina Los for library preparation and sequencing. We thank Suzy Strickler for helpful discussions and Drew Larson for constructive criticism and helpful feedback on the manuscript.

## Data availability

The *Asclepias viridiflora* raw paired-end sequence reads are available on the NCBI Sequence Read Archive (SRA) under BioProject PRJNA1263517. Accession numbers for the publicly available raw paired-end sequence reads used in this study can be found in Supplementary Table 1. Assemblies, orthologs, homologs, species/gene tree files, gene/whole genome duplication files, and BUSCO analyses can be found on Zenodo (10.5281/zenodo.15652983).

## Supplementary Material

### Supplementary Table Captions

**Supplementary Table 1.** Samples used in this study, along with details regarding the sequence processing and the assembly.

**Supplementary Table 2.** Full results of the GO term analysis.

**Supplementary Table 3.** Credits for photos used in Figure 1.

### Supplementary Figure Captions

**Supplementary Figure 1.** The Basic Universal Single Copy Ortholog Assessment results. All species are ordered Alphabetically.

**Supplementary Figure 2.** Inferred species relationships based on 3366 orthologs. A) Tree inferred using the Maximum Quartet Support Species Tree approach, as implemented in Astral, branch lengths are in coalescent units. B) Tree inferred using ML based on the concatenate supermatrix approach, branch lengths are in subs/site.

**Supplementary Figure 3.** The ML inferred relationships among extracted chloroplast genes and mitochondrial genes. Branches are in subs/site and support values at the nodes are in ultrafast bootstraps.

**Supplementary Figure 4.** The inferred Ks plots for all species in the study.

**Supplementary Figure 5.** Relationships among the tribes sampled in this study, as inferred by previous work on the backbone of Apocynaceae.

**Supplementary Figure 6.** Species relationship with taxa that did not have a copy of “*S-locus lectin protein kinase family protein*” recovered highlighted in red.

