## Supplementary figures and images for "Going green: Recycling transcriptomes to infer evolutionary relationships, gene duplication, gene tree conflict, and patterns of molecular evolution in the Apocynaceae"

### Supplementary Figure 1

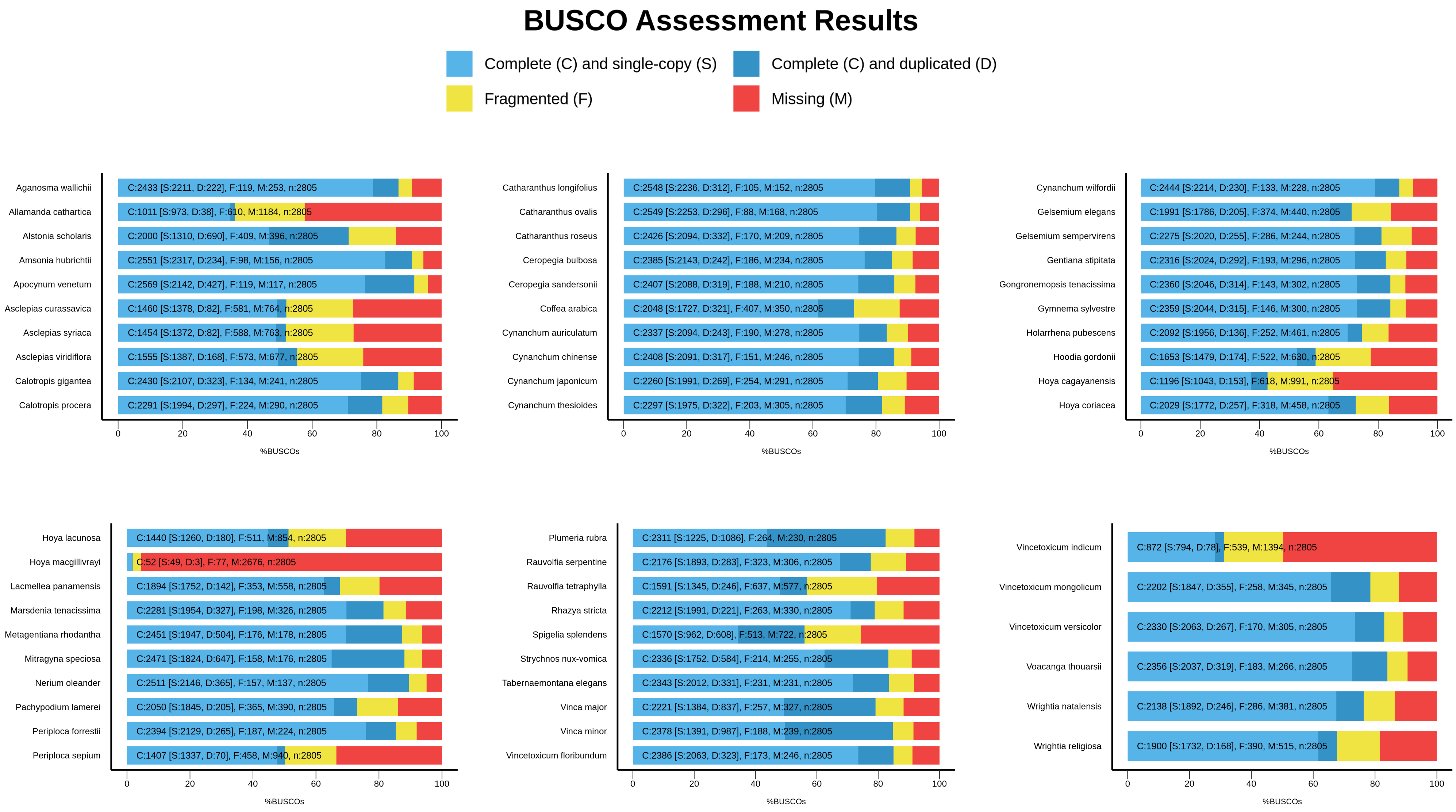

### Supplementary Figure 2

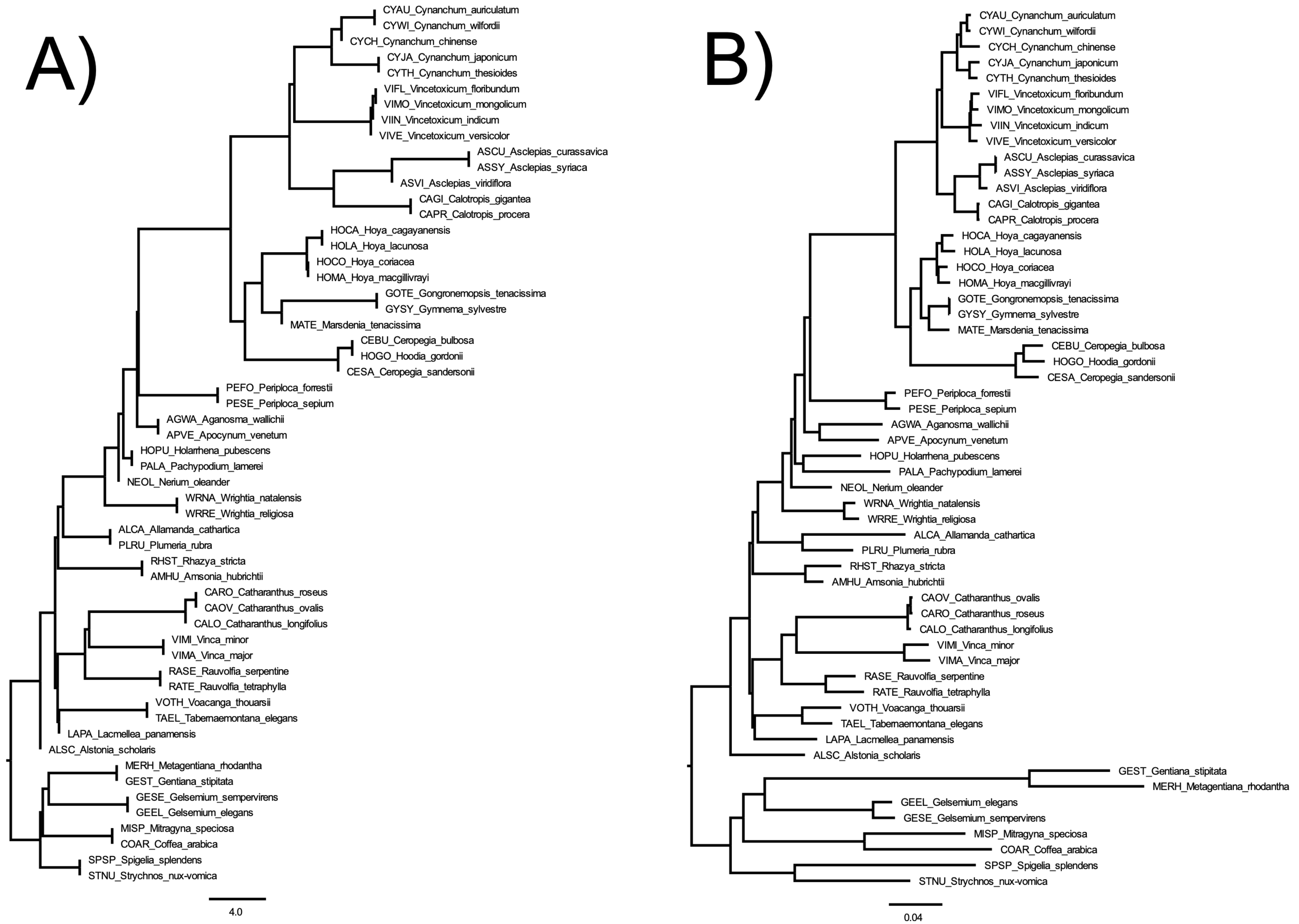

### Supplementary Figure 3

#### Chloroplast Inferred Phylogeny

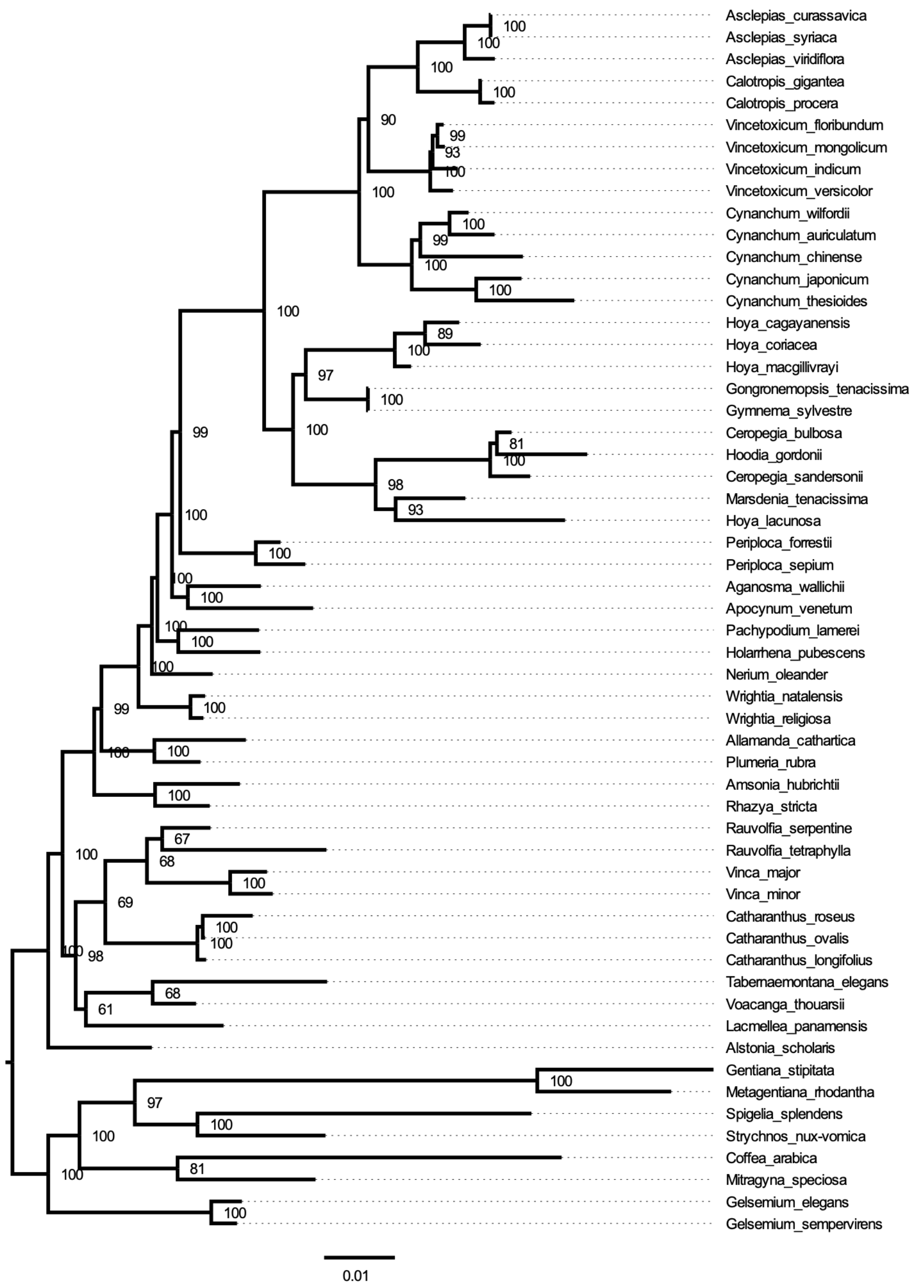


#### Mitochondria Inferred Phylogeny

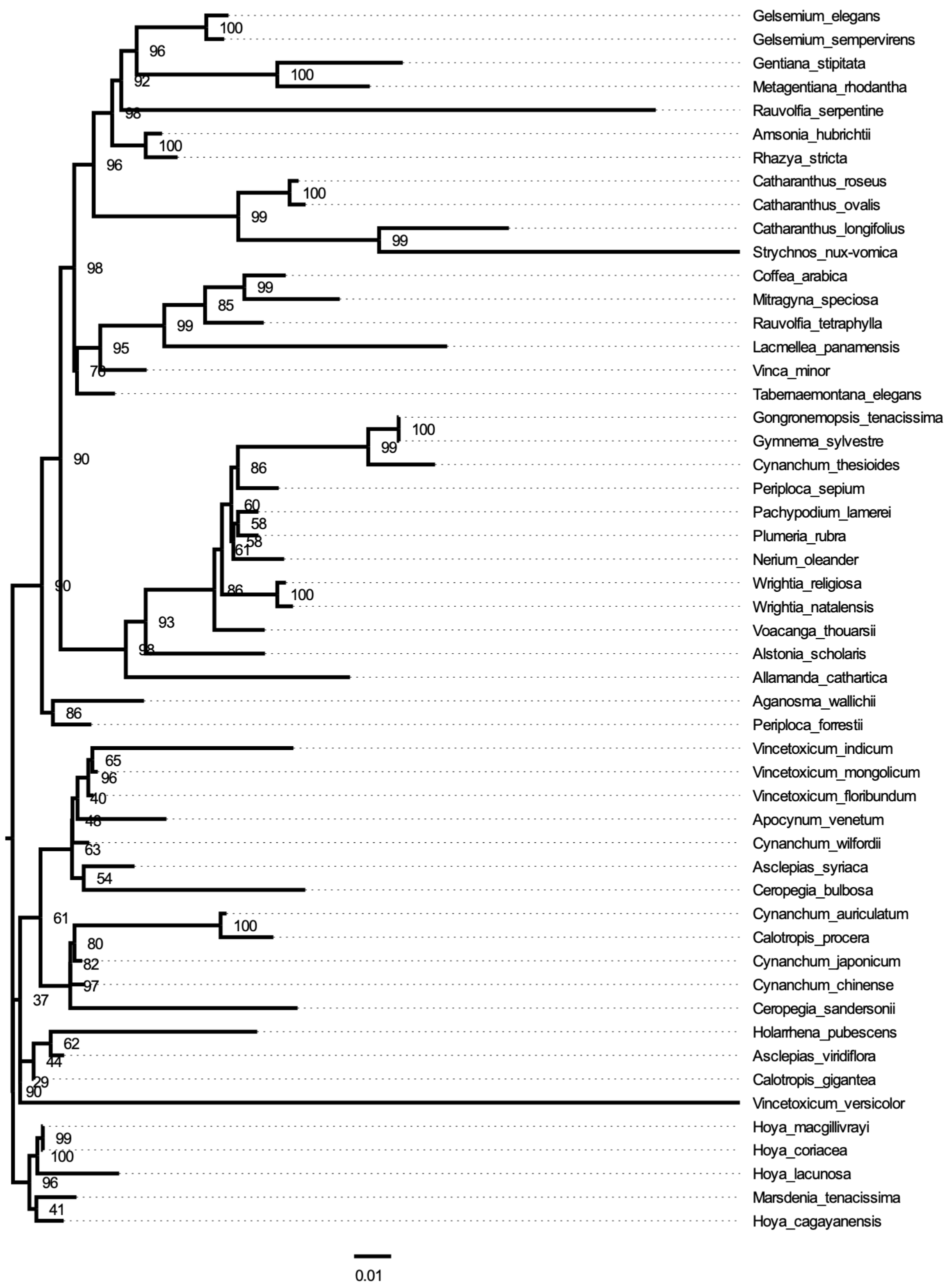

### Supplementary Figure 4

##### Outgroups

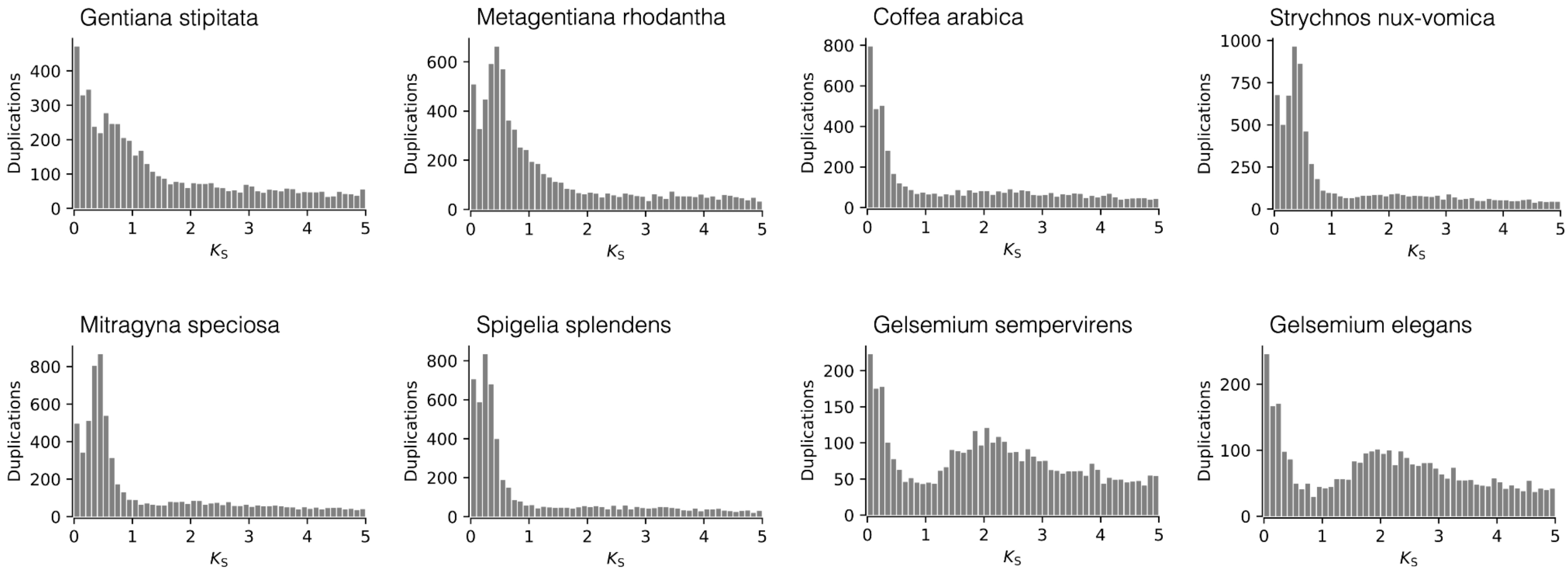


##### Ingroups

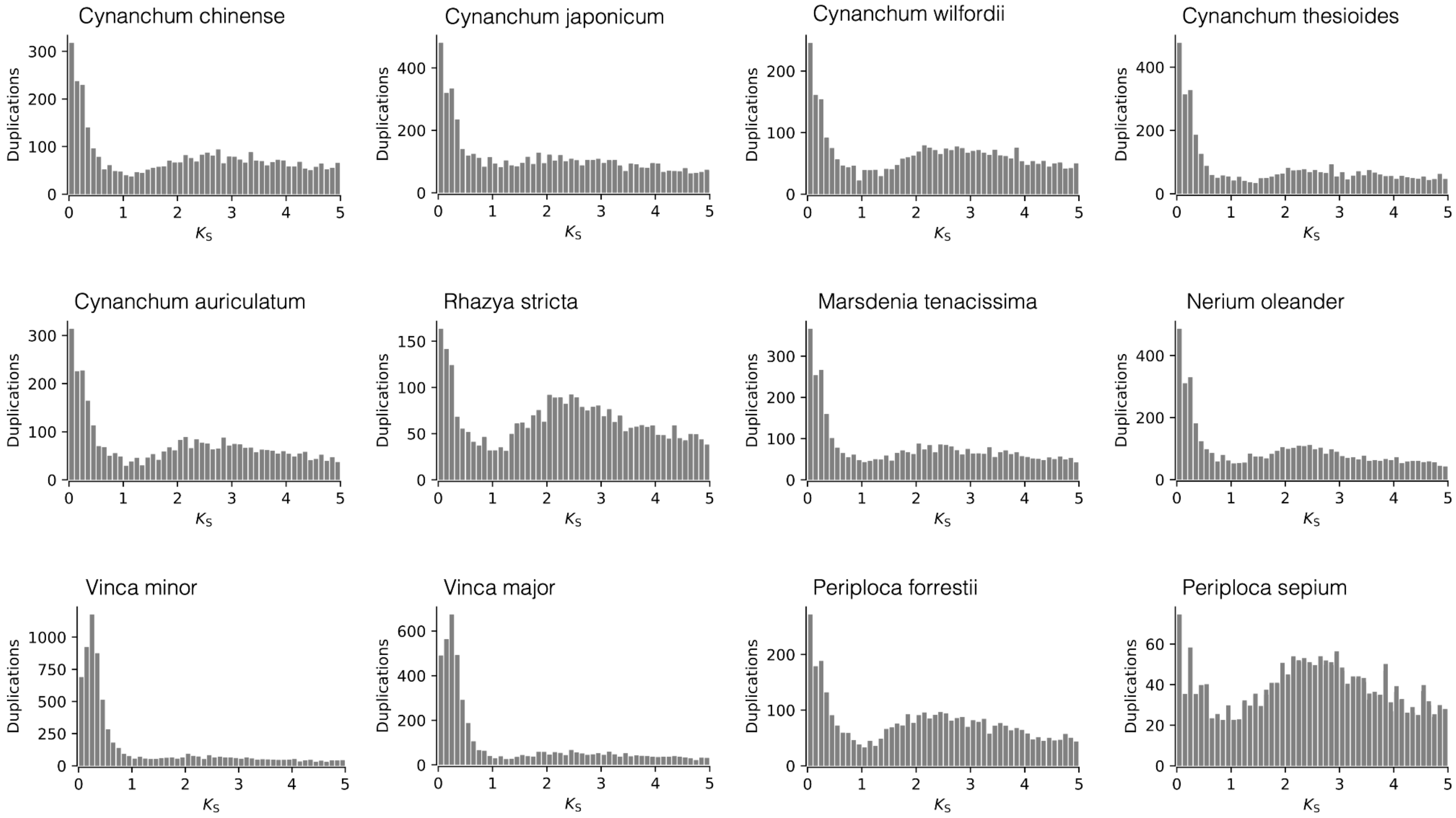

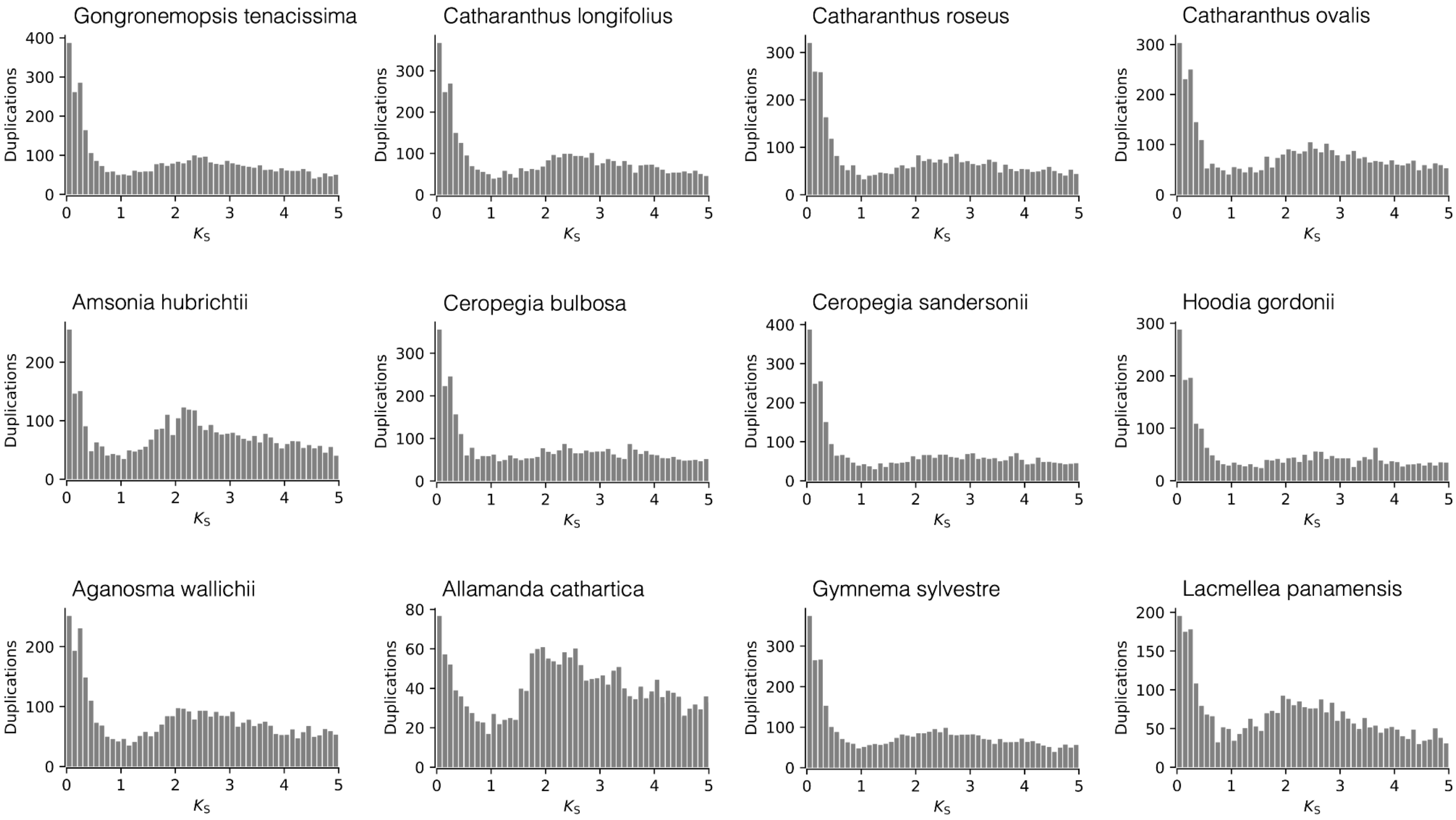

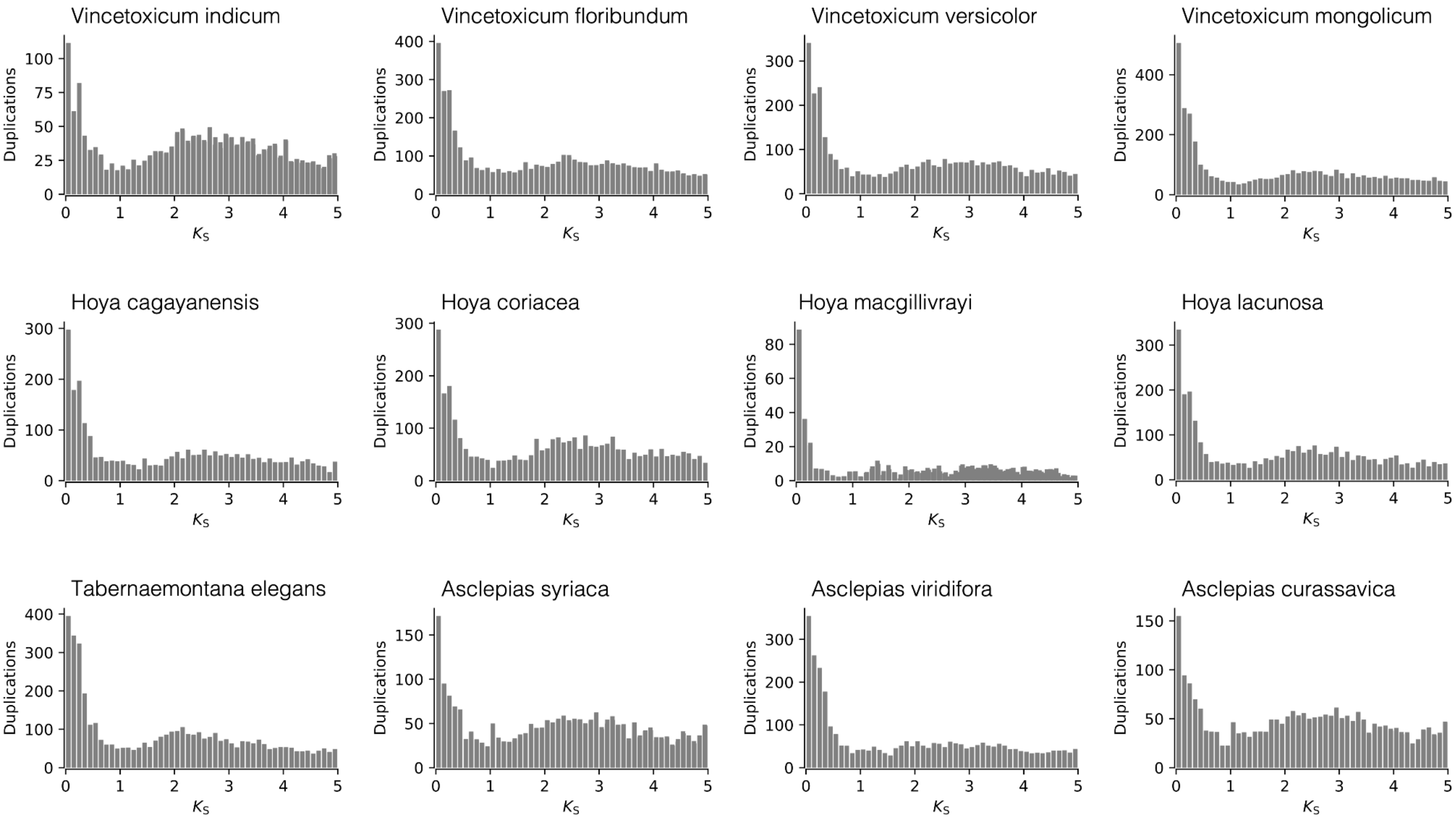

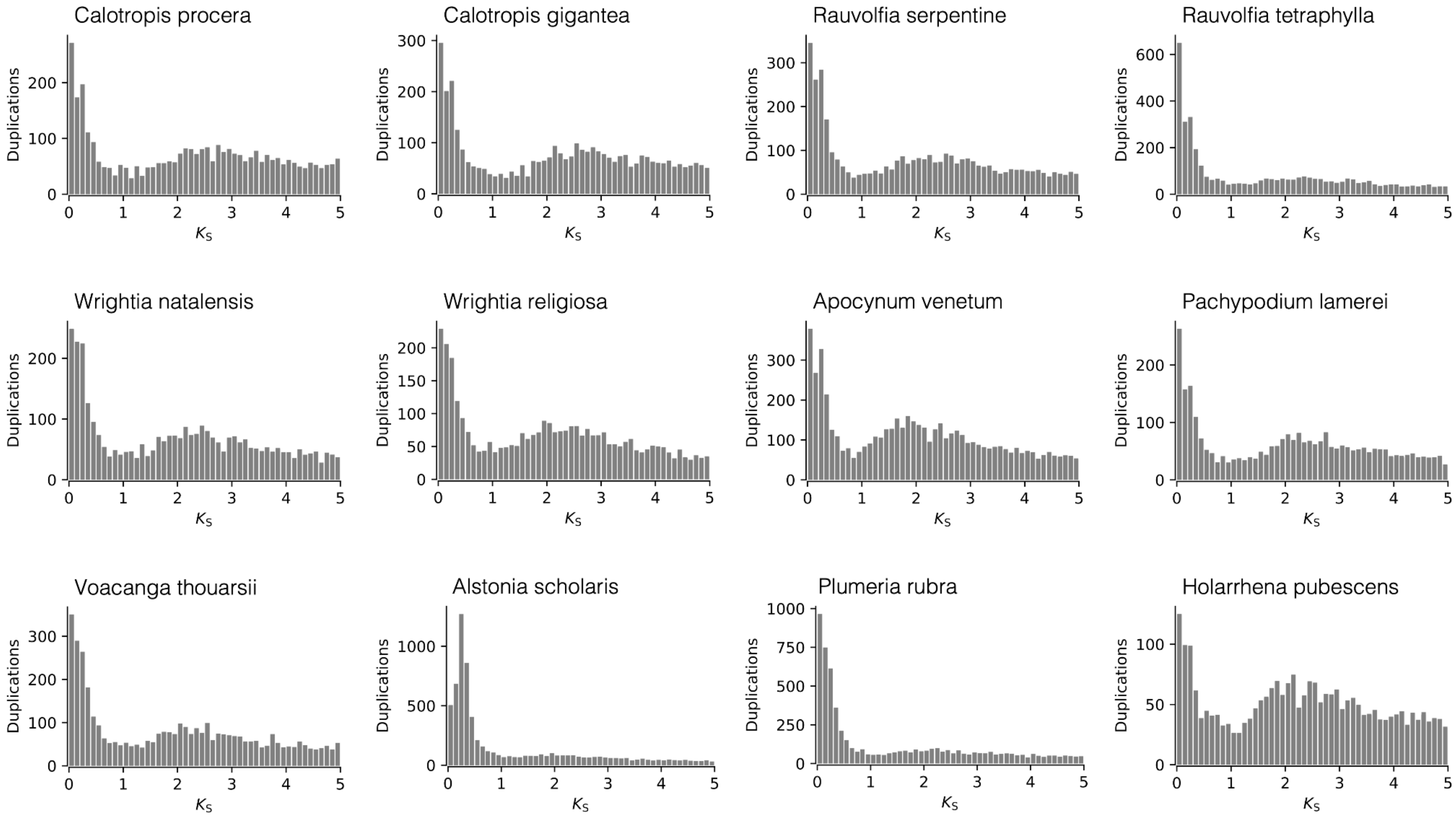

### Supplementary Figure 5

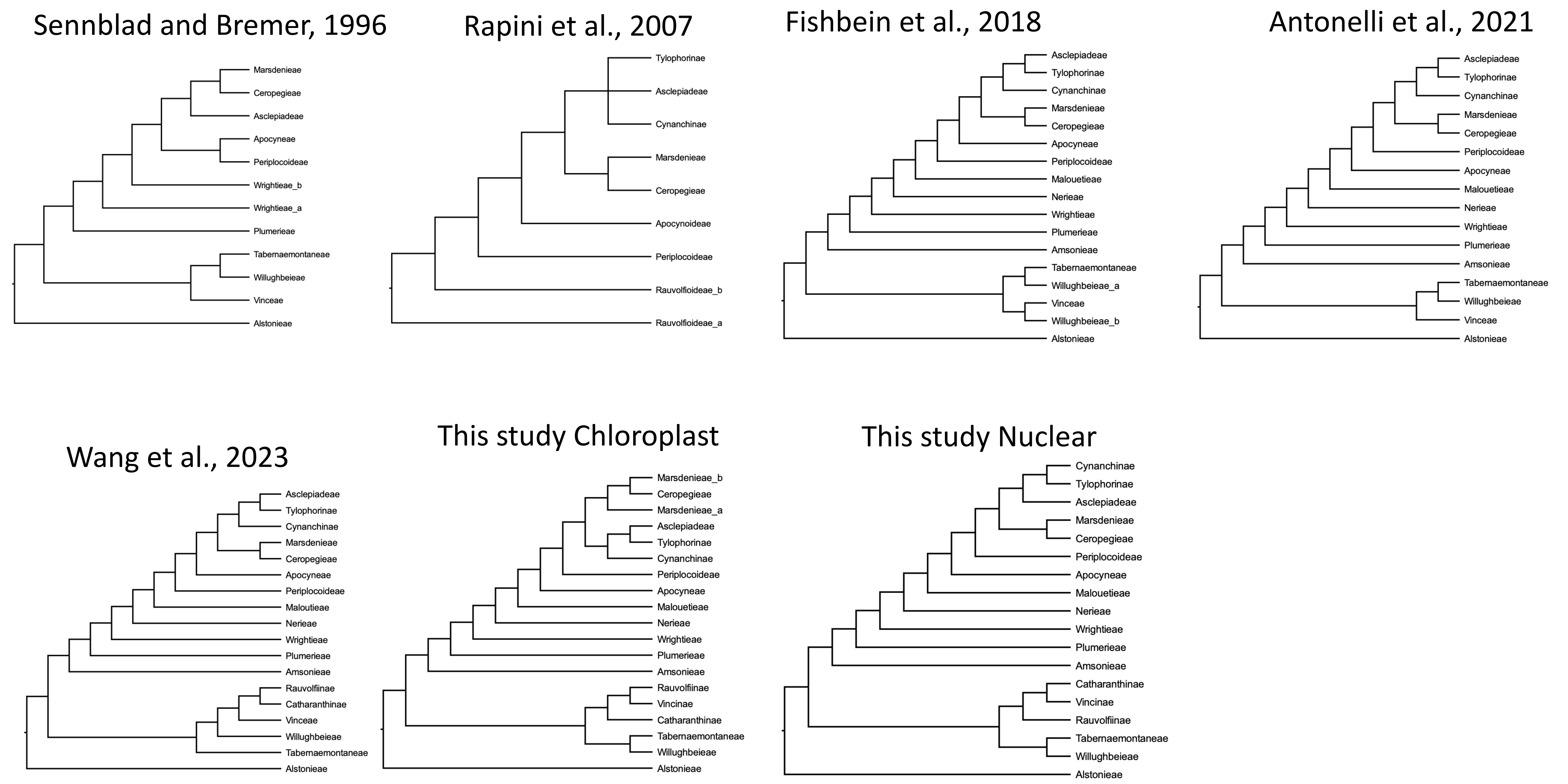

### Supplementary Figure 6

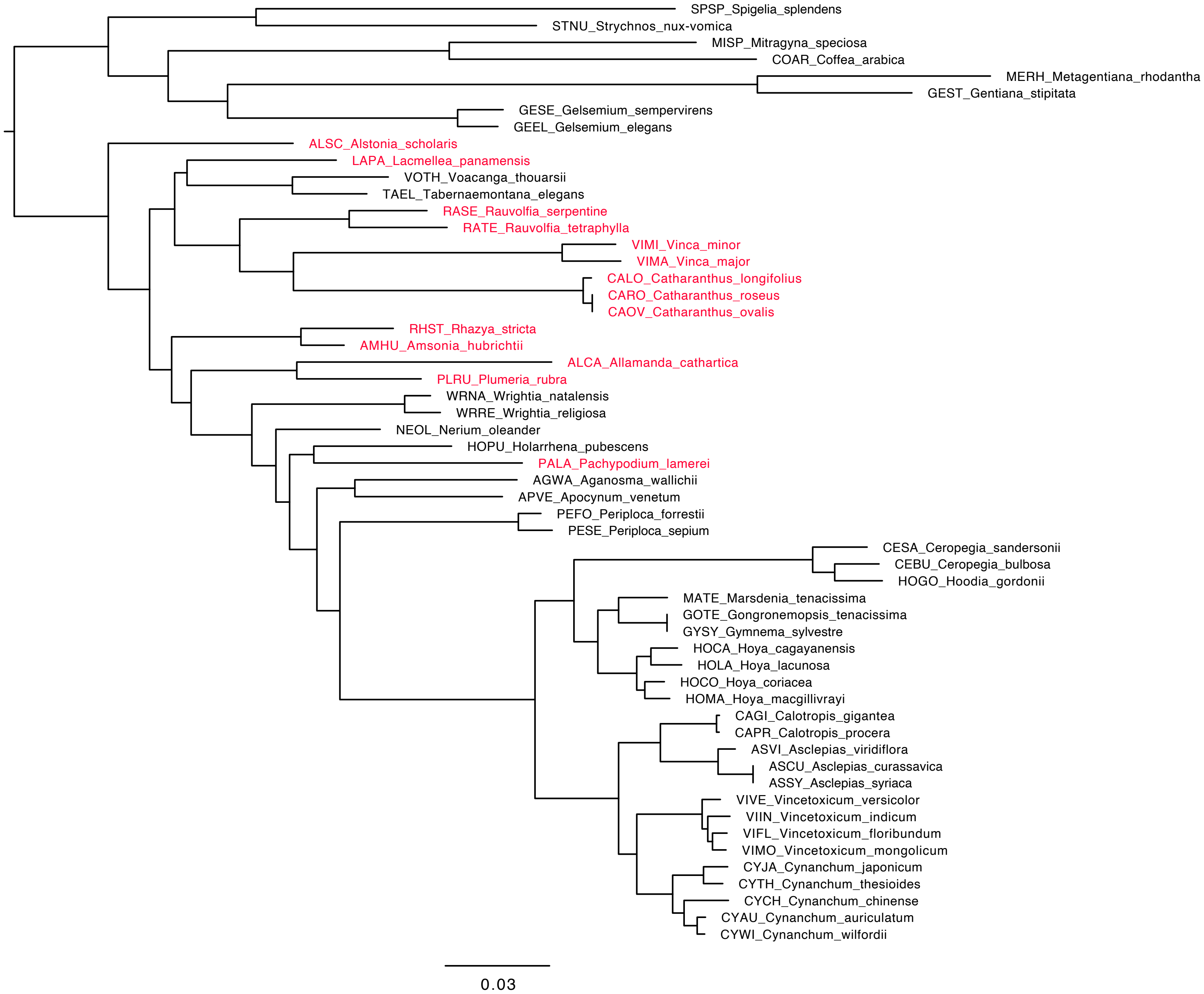
